# PCSK9 dependent cholesterol acquisition and utilization underlies metastatic organ preference in pancreatic cancer

**DOI:** 10.1101/2025.04.22.649912

**Authors:** Gilles Rademaker, Grace A. Hernandez, Sumena Dahal, Lisa Miller-Phillips, Alexander L. Li, Changfei Luan, Longhui Qui, Maude A. Liegeois, Bruce Wang, Kwun W. Wen, Grace E. Kim, Eric A. Collisson, Stephan F. Kruger, Stefan Boeck, Steffen Ormanns, Michael Guenther, Volker Heinemann, Michael Haas, Jen Jen Yeh, Roberto Zoncu, Rushika M. Perera

## Abstract

To grow at distant sites, metastatic cells must overcome major challenges posed by the unique cellular and metabolic composition of secondary organs^1^. Pancreatic ductal adenocarcinoma (PDAC) is an aggressive disease that metastasizes to the liver and lungs. Despite evidence of metabolic reprogramming away from the primary site, the key drivers that dictate the ability of PDAC cells to colonize the liver or lungs and survive there are undefined. We identified Proprotein Convertase Subtilisin/Kexin Type 9 (PCSK9) as predictive of colonization of the liver versus lungs by integrating datasets describing the metastatic tropism of human PDAC cell lines^2^, with in vivo metastasis modeling in mice and gene expression correlation analysis. PCSK9 is a negative regulator of low density lipoprotein(LDL)-cholesterol import and, accordingly, PCSK9-low PDAC cells exhibit strong preference for colonizing LDL-rich liver tissue. Cholesterol taken up by liveravid PCSK9-low cells is converted into the signaling oxysterol, 24(S)-hydroxycholesterol, which reprograms the surrounding microenvironment to induce nutrient release from neighboring hepatocytes. Conversely, PCSK9-high, lung-avid PDAC cells rely on transcriptional upregulation of the distal cholesterol synthesis pathway to generate intermediates, chiefly, 7dehydrodesmosterol and 7-dehydrocholesterol, with protective action against ferroptosis, a vulnerability in the oxygen-rich microenvironment of the lung. Increasing PCSK9 levels redirected liver-avid cells to the lung whereas its ablation drove lung-avid cells to the liver, thereby establishing PCSK9 as necessary and sufficient for secondary organ site preference. Our studies reveal PCSK9-driven differential utilization of the distal cholesterol synthesis pathway as a key and potentially actionable driver of metastatic growth in PDAC.

## Main text

Metastatic colonization requires infiltrating tumor cells to adapt to microenvironments that differ widely from their primary site of origin^3^. Our understanding of how this complex process occurs is limited, and the mechanisms that enable efficient outgrowth at secondary organ sites characterized by different nutrient, cellular and structural features, remain unclear. Metabolic plasticity of tumor cells is a major determinant of survival in circulation and subsequent adaptation and colonization within new tissue niches^1^. Likewise, specific metabolic features of the destination organ exert strong selective pressure on incoming tumor cells, such that only those with favorable pre-existing or evolved traits will ultimately survive and give rise to the colonizing population^4^. PDAC is an aggressive malignancy that commonly metastasizes to the liver, lungs, peritoneum, and lymph nodes^5^ however the specific target organ has relevance for disease progression as patients presenting with lung restricted metastases have a more favorable prognosis than those with liver or multi-site metastases^5–9^. Several pathways and processes have been linked to PDAC metastasis^10–16^ however we have an incomplete picture of what determines target organ preference, and what factors enable tumor cells to grow once they reach a specific organ site. Here we show that differential utilization of the distal cholesterol biosynthesis pathway is a major determinant of the ability of PDAC cells to colonize the liver versus the lung. Through its ability to control cholesterol import, the Proprotein Convertase Subtilisin/Kexin Type 9 (PCSK9) emerges as a central regulator of the availability of specific cholesterol intermediates and derivatives, which in turn play key roles during adaptation of PDAC cells to niche specific challenges within the liver and lung microenvironments.

### Human PDAC cell lines display differential preference for colonization of the liver and lungs

Recent development of high throughout pooled *in vivo* screens utilizing bar-coded human cancer cell lines^2^ revealed organ-specific patterns of metastasis across tumor types of diverse lineages, including pancreatic ductal adenocarcinoma (PDAC) – an aggressive malignancy that commonly metastasizes to the liver and lungs. To stratify PDAC cells according to organ colonization preference, we utilized these screening data (https://depmap.org/metmap/) to derive a fractional seeding ratio for 25 PDAC cell lines across the five assessed metastatic organ sites (liver, lung, kidney, brain, bone) (**Fig. 1a,b**; **Extended Data Fig. 1a,b**). Our analysis identified two subsets of PDAC – one showing clear avidity for growth in the liver (cluster 1) and another displaying a preference for growth in other organs, including the lungs (cluster 2) (**Fig. 1c**). Cluster 1 lines had a calculated mean metastatic potential of 70.6% and penetrance of 37.9% for colonization of the liver (**Supplementary Table 1**). In contrast, cluster 2 lines showed low metastatic potential and penetrance towards the liver (3.5% and 9.8% respectively) and a greater avidity towards other organs such as the lung (31.2% metastatic potential and 51.3% metastatic penetrance) (**Extended Data Fig. 1a,b; Supplementary Table 1**).

**Figure 1.**
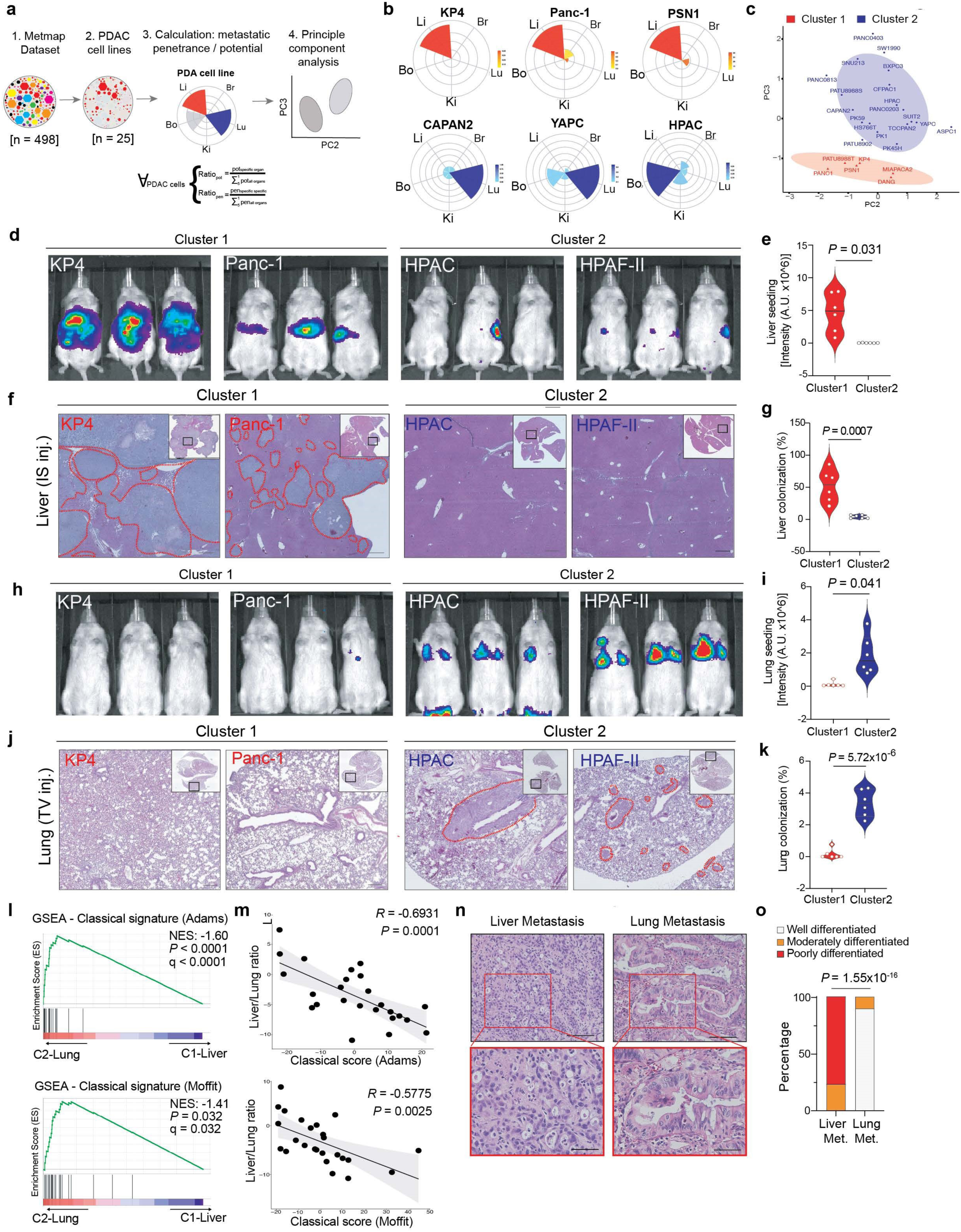
**a.** Workflow for deriving metastatic potential and penetrance of PDAC cell lines using the MetMap500 database. **b.** Petal plots depicting metastatic potential (length) and penetrance (colour) for the indicated cell lines. Br, Brain; Bo, Bone; Ki, Kidney; Li, Liver; Lu, Lung. **c.** Scatter plot representing two principal components (PC2 and PC3) derived from PCA analysis of PDAC metastatic potential and penetrance data. Each point represents an individual PDAC cell line. Two clusters are depicted. **d.** Normalized bioluminescence imaging following intra-splenic injection of NSG mice with the indicated cluster 1 and cluster 2 cell lines (n=6 mice/cluster) stably expressing luciferase. **e.** Cumulative quantification of bioluminescence signal from cluster 1 and 2 cell lines. **f.** Haematoxylin & Eosin (H&E) staining showing tumour invasion in the liver, with lesions outlined in red. Scale bars: 500 µm. **g.** Quantification of percentage tumour area calculated from images in f. **h.** Normalized bioluminescence imaging following tail-vein injection of NSG mice with the indicated cluster 1 and cluster 2 cell lines (n=6 mice/cluster) stably expressing luciferase. **i.** Cumulative quantification of bioluminescence signal from cluster 1 and 2 cell lines. **j.** H&E staining showing tumour invasion in the lungs, with lesions outlined in red. Scale bars: 500 µm. **k.** Quantification of percentage tumour area calculated from images in j. **l.** Gene set enrichment analysis (GSEA) plots showing relative enrichment of the PDAC classical gene signature derived from Adams et al. (top) and Moffit et al. (bottom) in cluster 1 (C1-Liver) and cluster 2 (C2-Lung) cell lines. **m.** Scatter plot showing correlation between liver/lung metastatic potential ratio (see Supplementary Table 2) and classical score from Adams et al.(top) and Moffit et al. (bottom). Each dot represents a PDAC cell line (n=25). **n.** H&E staining of human liver and lung PDAC metastases. Red boxes represent insets shown below. **o.** Bar plot showing the differentiation status of PDAC liver and lung metastasis shown in n, (Liver metastases n=45; Lung metastases n=10).

To validate the metastatic potential of the predicted PDAC clusters, we selected 4 human PDAC cell lines (KP4, Panc-1, HPAC and HPAF-II) for individual characterization by assessing their ability to colonize the liver and lungs following intrasplenic (IS) and tail vein (TV) injection, respectively. Intrasplenic injection of 500,000 KP4 or Panc-1 cells led to tumor formation in the liver 4 weeks post-injection, while HPAC or HPAF-II cells did not form tumors in the liver during the same time frame (**Fig. 1d-g**). In contrast, tail vein injection of KP4 or Panc-1 cells did not lead to overt or histological evidence of tumor growth in the lungs, however, HPAC or HPAF-II cells avidly formed tumors in the lungs (**Fig. 1h-k**). To further validate these findings, each cell line was introduced into the blood stream via intracardiac injection to allow colonization of any organ site. Consistent with the intrasplenic and tail vein injection experiments, KP4 and Panc-1 cells avidly colonized the liver and not the lungs, while HPAC and HPAF-II cells displayed the opposite phenotype, showing preferential growth in the lungs and not the liver (**Extended Data Fig. 1c-d**). Of note, all 4 cell lines harbor identical activating *KRAS* mutations (Glycine 12 to Aspartate) and inactivation of *CDKN2A* and *TP53* tumor suppressor genes, indicating that genetic drivers of PDAC likely do not determine secondary colonization preference. Taken together, these results suggest that secondary organ preference can be modeled *in vivo* using human PDAC cell lines, and we identify two subsets displaying avidity towards the liver (cluster 1: C1-Liver) or lungs (cluster 2: C2-Lung).

### Liver versus lung colonization correlates with PDAC subtype

Prior transcriptional profiling studies have shown that PDAC can be stratified into two main transcriptional subtypes termed Basal and Classical – which are linked to distinct molecular features and patient prognosis^17–21^. For instance, basal PDAC display features of poor differentiation while classical PDAC are well differentiated. We find that classical PDAC gene signatures were enriched in C2-Lung lines (**Fig. 1l**), while liver avidity was anti-correlated with the classical gene signature (**Fig. 1m; Supplementary Table 2).** Accordingly, histological evaluation of human PDAC metastatic lesions showed that most lung metastases (N=9/10) were well differentiated and displayed higher nuclear staining intensity for GATA6 – a transcription factor strongly associated with the classical subtype^22^ – relative to liver metastases (**Extended Data Fig. 1e-f**). In contrast, liver metastases (N=35/45) display characteristics of poor differentiation (**Fig. 1n,o; Supplementary Table 3**). These results are consistent with prior studies showing that liver metastases tend to show features of poor differentiation found in basal PDAC cells^20,22–24^.

### Differential PCSK9 expression correlates with lung versus liver avidity

To determine molecular features that distinguish liver versus lung avid lines, we utilized gene expression correlation (https://depmap.org/) to identify transcripts specifically enriched in the GATA6-positive lung-avid PDAC lines. This analysis identified PCSK9 as the top ranked gene transcript correlating with high GATA6 expression (**Fig. 2a**). High levels of *PCSK9* transcript and protein were positively correlated with additional classical subtype associated genes including *CDH1* (encoding E-cadherin) (**Fig. 2b; Extended Data Fig. 2a,b**), *S100P* and *FOXQ1* (**Extended Data Fig. 2a,b**) and anti-correlated with basal subtype markers, including *ZEB1* (**Figure 2b; Extended Data Fig. 2a,b**) and *VIM* (**Extended Data Fig. 2a,b**). PCSK9 is a serine protease that modulates cholesterol homeostasis by binding to the low-density lipoprotein (LDL) receptor (LDLR) and promoting its lysosomal degradation, thereby suppressing LDL import (**Fig. 2c**). Accordingly, cells expressing high levels of PCSK9 mRNA showed increased secretion of PCSK9 protein (**Extended Data Fig. 2c**), as well as reduced total (**Fig. 2b**) and cell surface levels of LDLR protein (**Extended Data Fig. 2d,e**). In contrast, PCSK9 expression was barely detectable or absent in C1Liver cell lines (**Fig. 2b**), which instead expressed high levels of total and cell surface LDLR protein (**Fig. 2b; Extended Data Fig. 2d,e**). Similarly, mouse cell lines derived from a genetically engineered mouse model of PDAC (referred to as KPC^25^), also displayed differential PCSK9/LDLR levels, with a clear anti-correlation between these two proteins (**Extended Data Fig. 2f**). Moreover, PCSK9 low KPC lines showed avidity for growth in the liver, with some growth in the lung, while PCSK9 high lines efficiently colonized the lungs and not the liver (**Extended Data Fig. 2g,h**).

**Figure 2.**
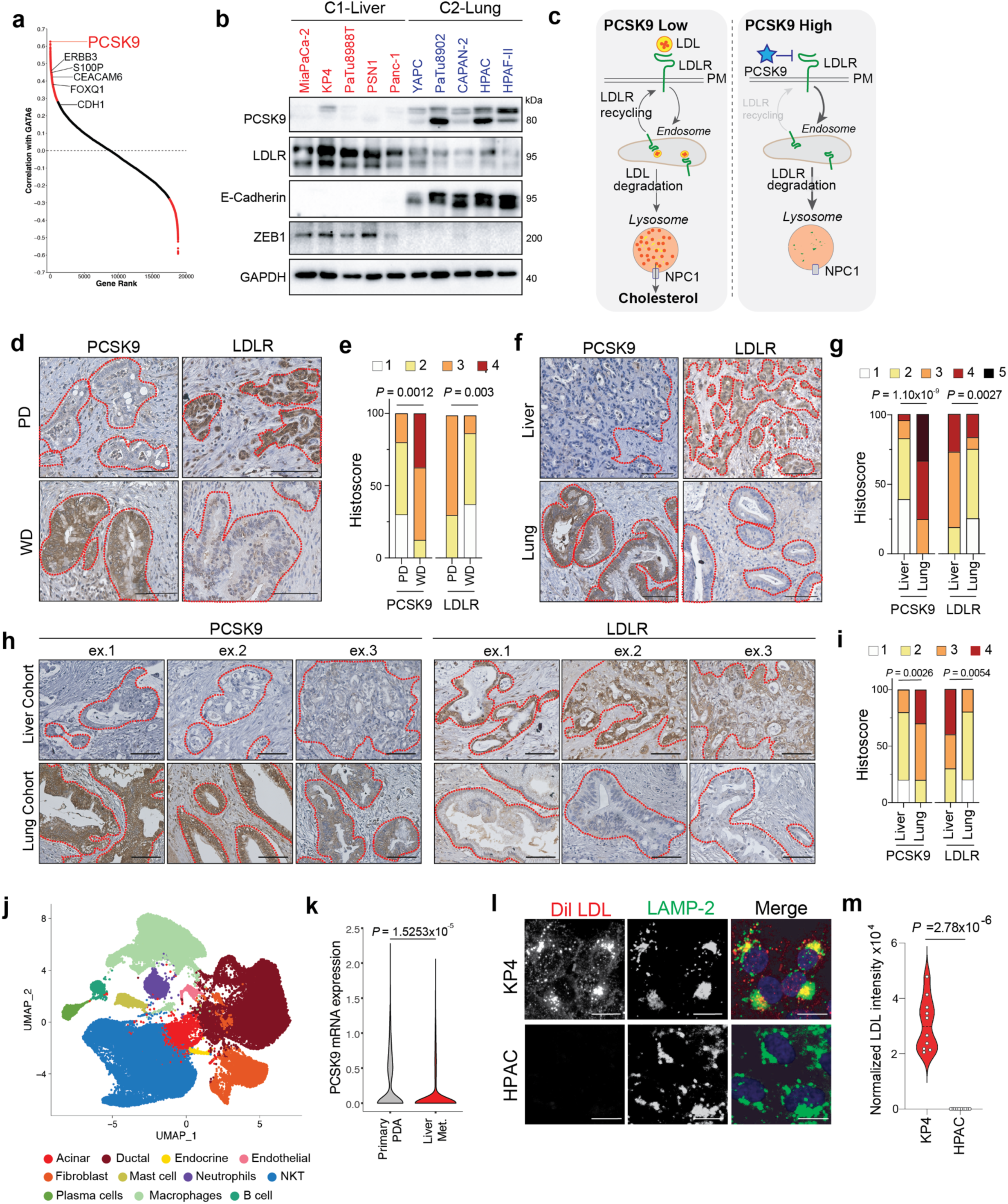
**a.** Analysis of top ranked gene expression correlations with *GATA6* showing *PCSK9* as the top correlated gene as well as other classical PDAC subtype genes. **b.** Western blot showing PCSK9 and LDLR levels and classical (E-cadherin) and basal (ZEB1) levels in C1-Liver and C2-Lung cell lines. **c.** Schematic of LDL uptake via LDL receptor (LDLR) and trafficking to the lysosome. High PCSK9 levels promotes LDLR endocytosis and lysosome degradation thereby inhibiting LDL uptake. **d.** Immuno-histochemical staining for PCSK9 and LDLR in poorly differentiated (PD) and well differentiated (WD) primary PDAC tumours. Tumour lesions are highlighted in red. Scale bars: 100 µm. (Poorly Differentiated n=10, Well Differentiated n=8). **e.** Bar graphs showing percentage positive intensity staining of samples in d. **f.** Immuno-histochemical staining for PCSK9 and LDLR in PDAC liver (n=26) and lung (n=12) metastases. Tumour lesions are highlighted in red. Scale bars: 100 µm. **g.** Bar graphs showing percentage positive intensity staining of samples in f. **h.** Immuno-histochemical staining for PCSK9 and LDLR in primary PDAC tissues from patients who developed liver-only metastasis (liver cohort) or lung-only metastasis (lung cohort). Scale bars: 100 µm (n=10/cohort). **i.** Bar graphs showing percentage positive intensity staining of samples in h. **j.** UMAP plot of single cell sequencing from primary PDAC (n=3) and matched liver metastases (n=4). **k.** PCSK9 mRNA expression in ductal cells corresponding to primary PDAC and liver metastases. **l.** Confocal microscopy images depicting Dil-LDL uptake (red) in C1-Liver (KP4) and C2-Lung (HPAC) cell lines after 20 minutes of incubation. Co-staining with LAMP2 (green) highlights lysosomal localization of the internalized LDL (yellow). Scale bars: 10 µm. **m.** Quantification of LDL intensity per field from images in l. (n=10 fields per samples)

To evaluate PCSK9 and LDLR expression in primary and metastatic PDAC, we first performed immunohistochemistry on primary PDAC patient specimens. Poorly differentiated (PD) primary PDAC and liver metastases displayed low PCSK9 and high LDLR levels, while well differentiated (WD) primary PDAC and lung metastases displayed high PCSK9 and low LDLR levels (**Fig. 2d-g; Supplementary Table 4,5**). We next utilized a matched cohort of primary PDAC from patients who developed liver only (liver cohort) or lung only (lung cohort) metastases (**Fig. 2h,i; Supplementary Table 6**). Liver cohort PDAC tumors characteristically showed low PCSK9 and high LDLR staining intensity within the ductal epithelia, whereas lung cohort PDAC tumors showed the opposite phenotype (eg. high PCSK9 and low LDLR staining). To further evaluate PCSK9 status in human PDAC, we analyzed single-cell RNA sequencing data from primary PDAC and the matched liver metastases^26^. Consistent with our immunohistochemical analysis, *PCSK9* expression is decreased in liver metastases relative to the matched primary tumor (**Fig. 2j,k**) Together, these results indicate that the PCSK9-low status is correlated with poorly differentiated PDAC and liver metastases, whereas the PCSK9-high status is associated with well differentiated PDAC and lung metastases.

### PCSK9-low liver-avid lines require LDL-cholesterol uptake for growth

We next conducted a series of experiments in C1-Liver and C2-Lung human PDAC cell lines and mouse KPC lines to mechanistically evaluate the functional role of PCSK9 in linking regulation of cholesterol homeostasis to target organ selection and colonization. First, C1-Liver cells showed high LDL uptake relative to C2-Lung cells as measured by immunofluorescence staining of DiLLDL uptake and trafficking to the lysosome (**Fig. 2l,m**; **Extended Data Fig. 3a,b**). Similarly, uptake of exogenous LDL was high in mouse PDAC cell lines with low PCSK9 levels (KPC1, KPC2) and absent in PCSK9 high cell lines (KPC3, KPC4) (**Extended Data Fig. 3c,d**). Moreover, Filipin staining, which visualizes cellular cholesterol depots, revealed high levels of lysosomal cholesterol in C1-Liver lines consistent with high cholesterol uptake rates, whereas C2-Lung lines predominantly displayed plasma membrane cholesterol and minimal lysosomal staining (**Extended Data Fig. 3e-g**). The combination of low PCSK9, high LDL uptake and lysosomal cholesterol, as measured by filipin staining, was also evident in cells isolated from a KPC derived liver metastases, relative to the matched isogenic primary PDAC cells (**Extended Data Fig. 3h-l**). Consistent with its high uptake, LDL was necessary for sustaining viability of C1-Liver lines but not C2-Lung lines. Specifically, C1-Liver cells and PCSK9 low KPC cells showed a marked decrease in proliferation following 96hrs of culture in lipoprotein depleted serum (LPDS), while the growth of C2-Lung lines was unaffected (**Fig. 3a,b; Extended Data Fig. 4a**). Lipoprotein depletion from C1-Liver lines led to a compensatory upregulation of cholesterol biosynthesis pathway enzymes, indicating activation of SCAP-SREBP regulatory feedback^27,28^ (**Extended Data Fig. 4b,c**); however, these enzymes were not further elevated by LDL depletion in C2-Lung lines (**Extended Data Fig. 4d**). Confirming its key requirement, addition of exogenous LDL or cholesterol was sufficient to rescue growth of C1-Liver lines grown in LPDS, indicating that loss of LDL-cholesterol is the primary mediator of the observed growth impairment (**Extended Data Fig. 4e,f**). Similarly, knockout of LDLR was selectively detrimental to C1-Liver lines relative to C2-Lung lines (**Fig. 3c,d**) and impaired growth of KP4 cells in the liver (**Extended Data Fig. 4g,h**).

**Figure 3.**
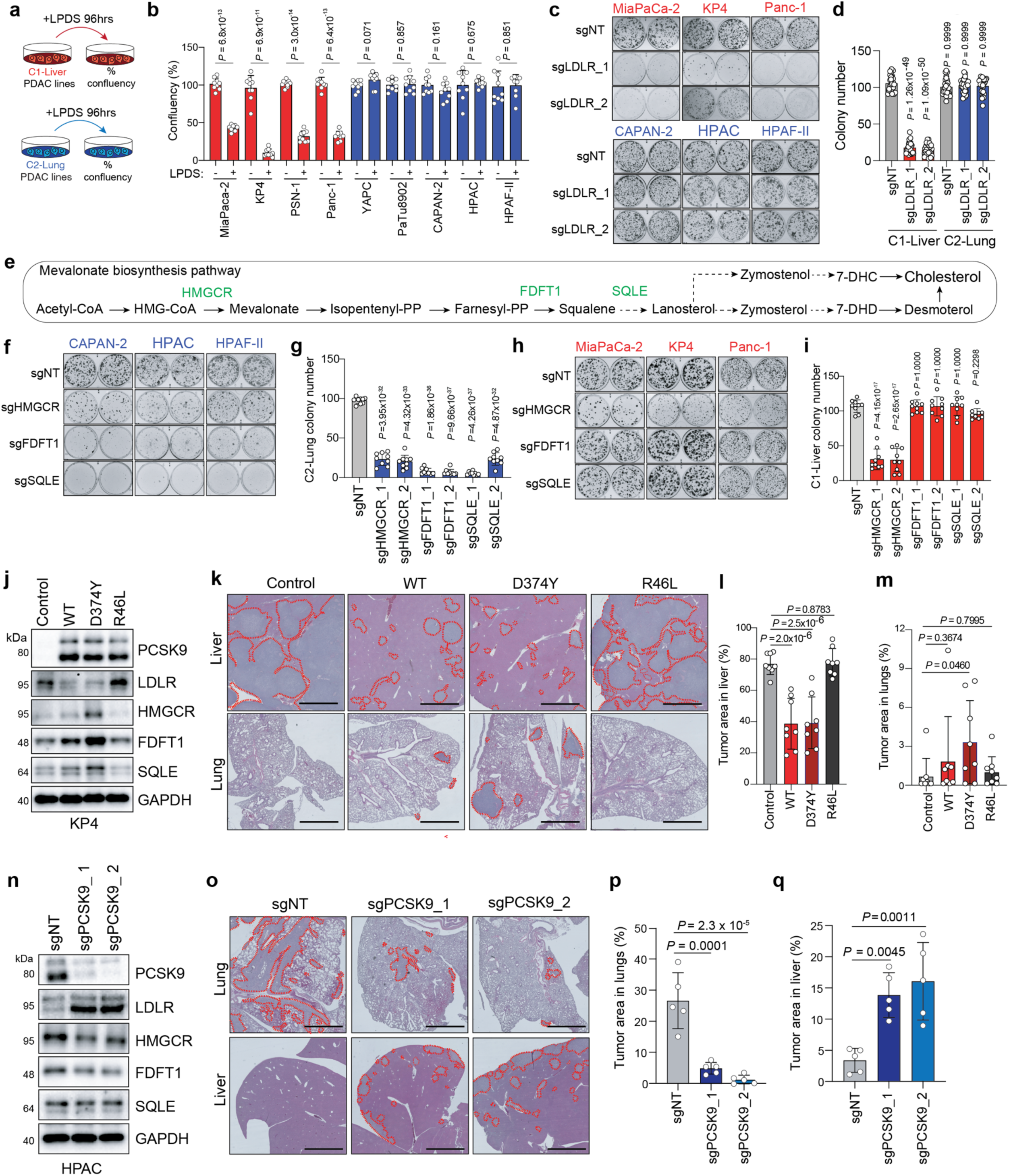
**a.** Schematic illustrating treatment of C1-Liver (KP4) and C2-Lung (HPAC) cells with lipoprotein depleted serum (LPDS) for 96hrs. **b.** Bar plot illustrating the percentage confluency of C1-Liver lines (red bars) and C2-lung lines (blue bars) following 96hrs of culture with LPDS. **c.** Colony forming ability of C1-Liver lines (MiaPaCa-2, KP4, and Panc-1) and C2-Lung lines (CAPAN2, HPAC, and HPAF-II) following CRISPR mediated knockout of LDLR. **d.** Quantification of colony forming number from images in c. **e.** Schematic illustrating reactions of the cholesterol biosynthesis pathway with key enzymes highlighted in green. **f.** Colony forming ability of the indicated C2-Lung lines following CRISPR-mediated knockout of the indicated cholesterol biosynthesis genes. **g.** Quantification of colony forming number from images in f. **h.** Colony forming ability of the indicated C1-Liver lines following CRISPR-mediated knockout of the indicated cholesterol biosynthesis genes. **i.** Quantification of colony forming number from images in h. **j.** Western blot of the indicated proteins in KP4 cells overexpressing PCSK9 variants. Wild type (WT), active variant (D374Y) and inactive variant (R46L). **k.** H&E staining of liver (top) and lungs (bottom) following intrasplenic or tail vein injection of KP4 cells harbouring the indicated PCSK9 variants, respectively into recipient mice. Tumor lesions are outlined in red. N=8 mice/condition. Scale Bar: 500µm **l.** Percentage tumour area in the liver from samples shown in k. **m.** Percentage tumour area in the lungs from samples shown in k. **n.** Western blot of the indicated proteins in HPAC cells following CRISPR mediated knockout for *PCSK9* with two independent sgRNAs. **o.** H&E staining of lungs (top) and liver (bottom) following tail vein or intrasplenic injection of HPAC cells following CRISPR mediated knockout of PCSK9, respectively into recipient mice. N=5 mice/conditions. Scale Bar: 500µm **p.** Percentage tumour area in the lungs from samples shown in o. **q.** Percentage tumour area in the liver from samples shown in o.

### PCSK9-high lung avid lines are dependent on cholesterol biosynthesis for growth

Consistent with the low exogenous LDL-cholesterol uptake in PCSK9-high C2-Lung lines and KPC lines, these cells displayed higher expression of genes responsible for cholesterol biosynthesis (**Extended Data Fig. 5a-c**). Moreover, analysis of DepMap gene dependency data showed that *PCSK9* expression in PDAC is positively associated with sensitivity to loss of late step cholesterol biosynthesis genes (**Extended Data Fig. 5d**). Consistent with this finding, knockout of distal cholesterol synthesis enzymes, Farnesyl-Diphosphate Farnesyltransferase 1 (FDFT1) and Squalene Epoxidase (SQLE), led to growth impairment of C2-Lung lines (**Fig. 3e-g; Extended Data Fig. 5e,f**), but had no effect on the growth of C1-Liver lines, likely due to their reliance on LDLR-mediated import of LDL-cholesterol (**Fig. 3h,i; Extended Data Fig. 5e,f**). Of note, all lines were sensitive to knockout of HMG-CoA reductase (HMGCR), which catalyzes the rate limiting step of the mevalonate pathway (**Figure 3e-i; Extended Data Fig. 5e,f**), suggesting that biosynthetic intermediates generated by the upper mevalonate pathway, such as farnesyl-pyrophosphate and its derived isoprenoids, are required for growth of all PDAC lines. This result is consistent with antitumor effects associated genetic deletion of *SCAP* in a PDAC mouse model^29^. Collectively, these results suggest that unlike C1-Liver lines, which obtain cholesterol via uptake, C2-Lung cell lines rely on de novo synthesis to maintain cholesterol pools and cellular growth.

### PCSK9 drives secondary organ selection

Given our findings that PCSK9 expression levels translated into differential dependency on exogenous cholesterol by the liver-versus lung-avid lines, we next tested whether enforced PCSK9 gain or loss of function is sufficient to drive preferential growth in the liver or lungs. First, the C1Liver line, KP4, was engineered to ectopically express either wild type (WT) PCSK9, a dominant active variant (D374Y) or a catalytically inactive variant (R46L)^30^ (**Fig. 3j**). Expression of WT and PCSK9^D374Y^ caused a decrease in LDLR levels (**Fig. 3j**), a corresponding decrease in LDL uptake (**Extended Data Fig. 6a,b**) and a shift towards *de novo* cholesterol biosynthesis, as evidenced by increased expression of mevalonate pathway enzymes (**Fig. 3j**). These stable cell lines were then transplanted into the liver or lungs and tumor growth was monitored over a period of 4 weeks. Parental KP4 cells developed large tumors within the liver, as did the cells expressing PCSK9^R46L^. In contrast, expression of either WT or PCSK9^D374Y^ significantly inhibited growth of KP4 cells in the liver (**Fig. 3k,l**). Strikingly, while the parental KP4 cells did not grow in the lungs following tail vein injection, expression of PCSK9^D374Y^ was sufficient to enable growth of KP4 cells in the lung (**Fig. 3k,m**).

To complement these experiments, we next tested whether knockout of PCSK9 in a C2-Lung line, HPAC, is sufficient to enable growth within the liver. CRISPR mediated knockout (KO) of PCSK9 in HPAC cells led to an increase in LDLR protein (**Fig. 3n**), increased LDL uptake (**Extended Data Fig. 6c,d**) and decreased expression of cholesterol biosynthesis genes (**Fig. 3n**), but had no effect on growth rate in vitro (**Extended Data Fig. 6e**). However, upon tail vein injection, PCSK9 KO cells grew less in the lungs and more in the liver, relative to the sgNT control HPAC cells (**Fig. 3o-q**). These findings support a model where PCSK9-LDLR status – and associated cholesterol uptake versus synthesis – is necessary and sufficient to drive growth of PDAC cells within the liver or the lungs, respectively.

Our findings show that baseline levels of PCSK9 predict secondary organ preference, especially in patients with liver- or lung-only metastases (see **Fig. 2h**). However, PCSK9-high and -low cells can coexist within the same primary tumor (**Extended Data Fig. 7a,b**). Therefore, we tested whether PCSK9 expression still predicts colonization in heterogenous tumors. Using mouse MT23 cells (shown in **Extended Data Fig. 3h**) – which form primary tumors with variable PCSK9 expression – we performed transplant injections either back into the pancreas via orthotopic transplantation, the liver via intrasplenic injection or the lungs via tail vein injection. After 2 weeks, mice were euthanized, tumors excised and immuno-stained for PCSK9 together with CK19 to identify pancreatic ductal epithelia. PCSK9 expression in the primary tumor was heterogeneous with some cells displaying high levels, while other cells had low expression (**Extended Data Fig. 7a,b).** In contrast, liver tumors displayed uniform downregulation of PCSK9 staining while lung tumors displayed uniformly high levels (**Extended Data Fig. 7a,b**). These results suggest that PCSK9 status reliably predicts organ-specific colonization and tumor burden in the liver or the lungs.

### Liver avid PDAC cells activate an oxysterol-mediated cholesterol efflux program in hepatocytes

We next sought to determine the fate of the LDL-cholesterol actively taken up by the C1-Liver cells, and its specific role in promoting survival and colonization of the liver. Cholesterol is a critical building block for cellular membranes and serves as a backbone for the synthesis of steroid hormones, bile acids and signaling metabolites such as oxysterols^27,28,31,32^. Quantitative RT-PCR showed that the mRNA level of *CYP46A1*, which encodes a cholesterol hydroxylase responsible for synthesizing 24(S)-hydroxycholesterol (24-OHC), was preferentially upregulated in C1-Liver cell lines relative to C2-Lung lines (**Fig. 4a**) and anti-correlated with PCSK9 expression in TCGA PDAC dataset (**Fig. 4b**). Of note, no other significant correlation between the expression of other cholesterol hydroxylases and PCSK9 status was observed (**Extended Data Fig. 8a,b**), indicating that CYP46A1 and its role in generation of 24-OHC synthesis may be important in C1-Liver cells.

**Figure 4.**
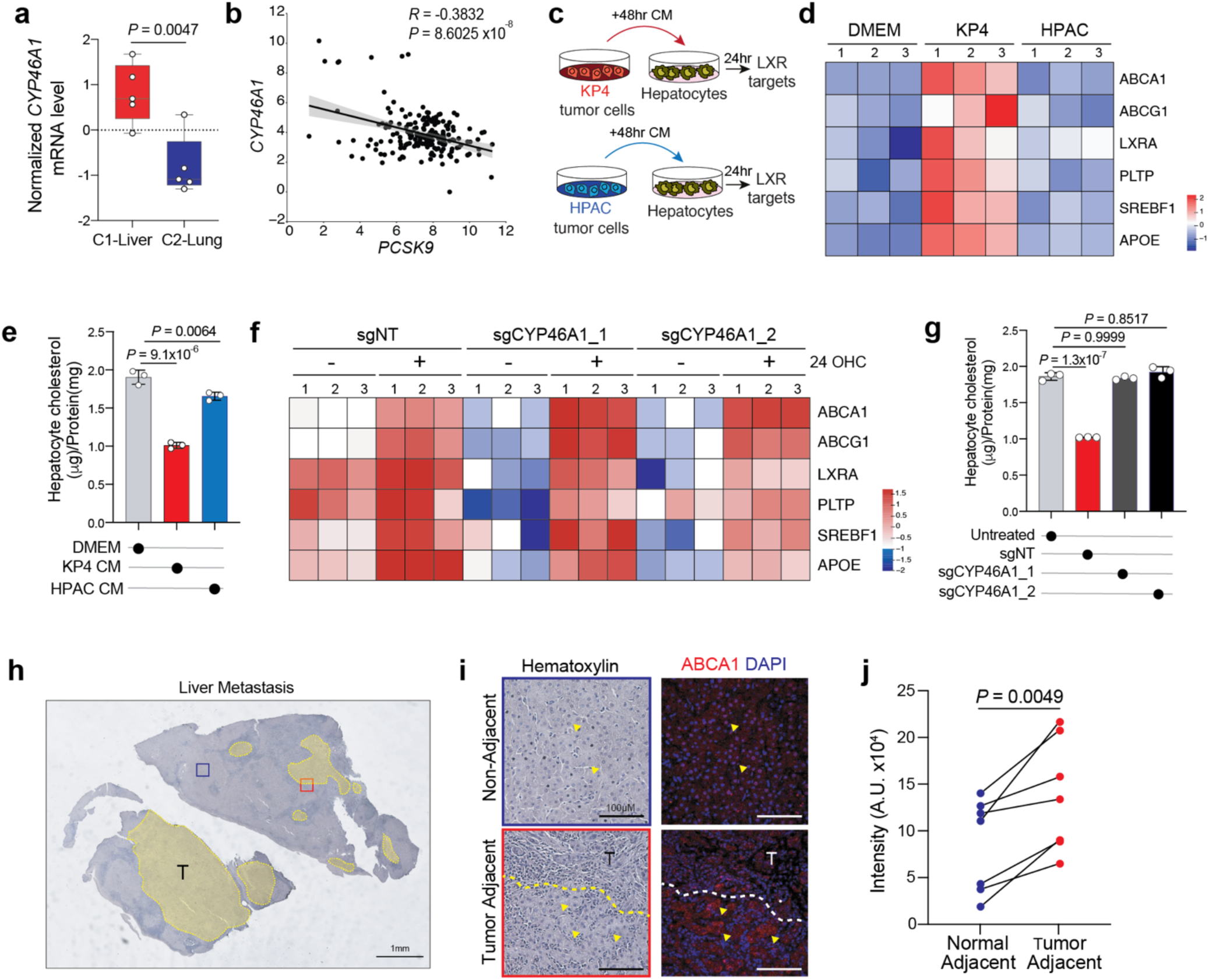
**a.** Normalized mRNA expression measured by quantitative real-time PCR of *CYP46A1* transcript in C1-Liver (KP4, MiaPaCa-2, Panc-1, PSN1, PaTu8988T) and C2-Lung (HPAC, HPAF-II, CAPAN2, YAPC and PaTu8902) lines. **b.** Anti-correlation between *PCSK9* and *CYP46A1* transcript expression in TGCA PAAD data. **c.** Schematic illustrating treatment of primary mouse hepatocytes with 48hr conditioned media (CM) isolated from C1-Liver (KP4) and C2-Lung (HPAC) cells. Hepatocytes were incubated for 24 hrs with CM and LXR target genes were subsequently measured by quantitative real-time PCR. **d.** Heatmap depicting the expression levels of LXR target genes in mouse hepatocytes following incubation with KP4 of HPAC CM for 24hrs. Each row represents a specific gene, while each threecolumn unit represents N=3 biological replicates per condition. Expression levels are expressed as z-scores, with positive z-scores indicating higher expression (red) relative to the mean and negative z-scores indicating lower expression (blue) relative to the mean. **e.** Total intracellular cholesterol in mouse hepatocytes following treatment with KP4 or HPAC CM for 24 hours, normalized to total cell number. N=3 biological replicates per condition. **f.** Heatmap depicting the expression levels of LXR target genes in mouse hepatocytes after treatment for 24hrs with CM isolated from KP4 parental and CYP46A1 knockout cells or treated with 24-hydroxycholesterol (24-OHC; 25µM). Each row represents a specific gene, while each three-column unit represents N=3 biological replicates per condition. Expression levels are expressed as z-scores, with positive z-scores indicating higher expression (red) relative to the mean and negative z-scores indicating lower expression (blue) relative to the mean. **g.** Total intracellular cholesterol in mouse hepatocytes following treatment with CM isolated from KP4 parental or CYP46A1 KO CM, normalized to cell number. N=3 biological replicates per condition. **h.** H&E image of a representative patient liver metastatic lesion. Yellow outlines indicate tumour regions. The red box indicates tumour adjacent region while the blue box represents a non-adjacent region. Scale bar: 1mm. **i.** H&E images (left) and immunofluorescence images of the cholesterol efflux transporter ABCA1 (right) of the regions indicated in h. Dashed line indicates the margin between tumor (T) and adjacent liver. Scale bar: 100µm. **j.** Quantification of ABCA1 staining intensity in tumor adjacent (red) and non-adjacent (blue) regions across N=7 liver metastases sections.

Studies in the brain have shown that 24-OHC, synthesized and secreted by neuronal cells, can promote activation of genes associated with lipoprotein synthesis and cholesterol efflux in neighboring astrocytes via the nuclear hormone receptor, Liver X Receptor (LXR)^33,34^. Given that hepatocytes are the primary source of LDL-cholesterol synthesis in the liver, we sought to determine whether an analogous tumor-hepatocyte circuit may also exist during colonization of the liver. To do so, LXR target gene activation was measured in primary mouse hepatocytes following incubation for 24hrs with conditioned media (CM) isolated from KP4 (C1-Liver) or HPAC (C2-Lung) cells (**Fig. 4c**). Only CM from KP4 cells activated LXR target genes (*ABCA1, ABCG1, LXRA, PLTP, APOE, SREBF1*) in mouse hepatocytes (**Fig. 4d**). Similarly, treatment with KP4 CM but not HPAC CM led to a decrease in hepatocyte cholesterol levels (**Fig. 4e**), suggesting an increased rate of hepatocyte cholesterol efflux occurs in response to KP4 CM.

The ability of KP4 CM to activate LXR target genes and promote cholesterol efflux by hepatocytes was dependent on CYP46A1 as *CYP46A1* ablation in KP4 cells completely blocked the ability of KP4 CM to induce LXR target genes and promote cholesterol efflux by hepatocytes (**Figure 4f,g**). Accordingly, addition of exogenous 24-OHC was able to rescue LXR target gene expression in hepatocytes treated with CYP46A1 KO KP4 CM (**Fig. 4f**). To assess whether LXR activation occurs in hepatocytes in vivo, tumor adjacent and non-adjacent sections from liver metastases were stained for the LXR transcriptional target ATP-binding cassette transporter 1 (ABCA1) (**Fig. 4h**), a cholesterol exporter belonging to the ABC family of transporters. We find that tumor adjacent hepatocytes express higher levels of ABCA1 relative to non-adjacent hepatocytes (**Fig. 4i,j**). This finding indicates that hepatocytes within proximity to tumor cells activate LXR target gene expression to a higher degree than non-adjacent hepatocytes. The most parsimonious rationale for the results is that C1-Liver derived 24-OHC serves to condition the liver microenvironment in a feed forward manner to continuously provide LDL-cholesterol for consumption by near-by tumor cells to sustain their growth.

### Lung avid PDAC cells produce distal cholesterol biosynthesis intermediates that protect against ferroptosis

Lastly, we sought to determine how the elevated de novo synthesis of cholesterol intermediates driven by high PCSK9 levels contributes to lung colonization by PDAC cells. Recent studies have shown that resistance to ferroptosis - an iron dependent form of cell death involving phospholipid peroxidation^35^, is mediated by several late step cholesterol synthesis intermediates including squalene^36^ and 7-dehydrocholesterol (7-DHC)^37–39^. The 5,7 diene B ring structure of 7-DHC functions as a radical-trapping agent that suppresses ferroptosis^37–39^, and sterols with similar structural features, such as 7-Dehydrodesmosterol (7-DHD), likely also have anti-ferroptotic capabilities (**Fig. 5a**). We find that high expression of Sterol C5-desaturase (SC5D), 7-DHC reductase (DHCR7) and DHCR24 expression, along with several other distal cholesterol biosynthesis pathway enzymes, are highly expressed and positively correlated with PCSK9-high status in PDAC patient samples (**Fig. 5b; Extended Data Fig. 9a**), and in C2-Lung lines relative to C1-Liver lines (see **Extended Data Fig. 5a,b**). Targeted metabolomics further showed higher 7DHD levels in HPAC cells relative to liver-avid KP4 cells (**Fig. 5c**) and ectopic expression of active PCSK9 variants (WT and/or D374Y) in KP4 cells was sufficient to increase the levels of 7-DHC and 7-DHD (**Fig. 5c,d**).

**Figure 5.**
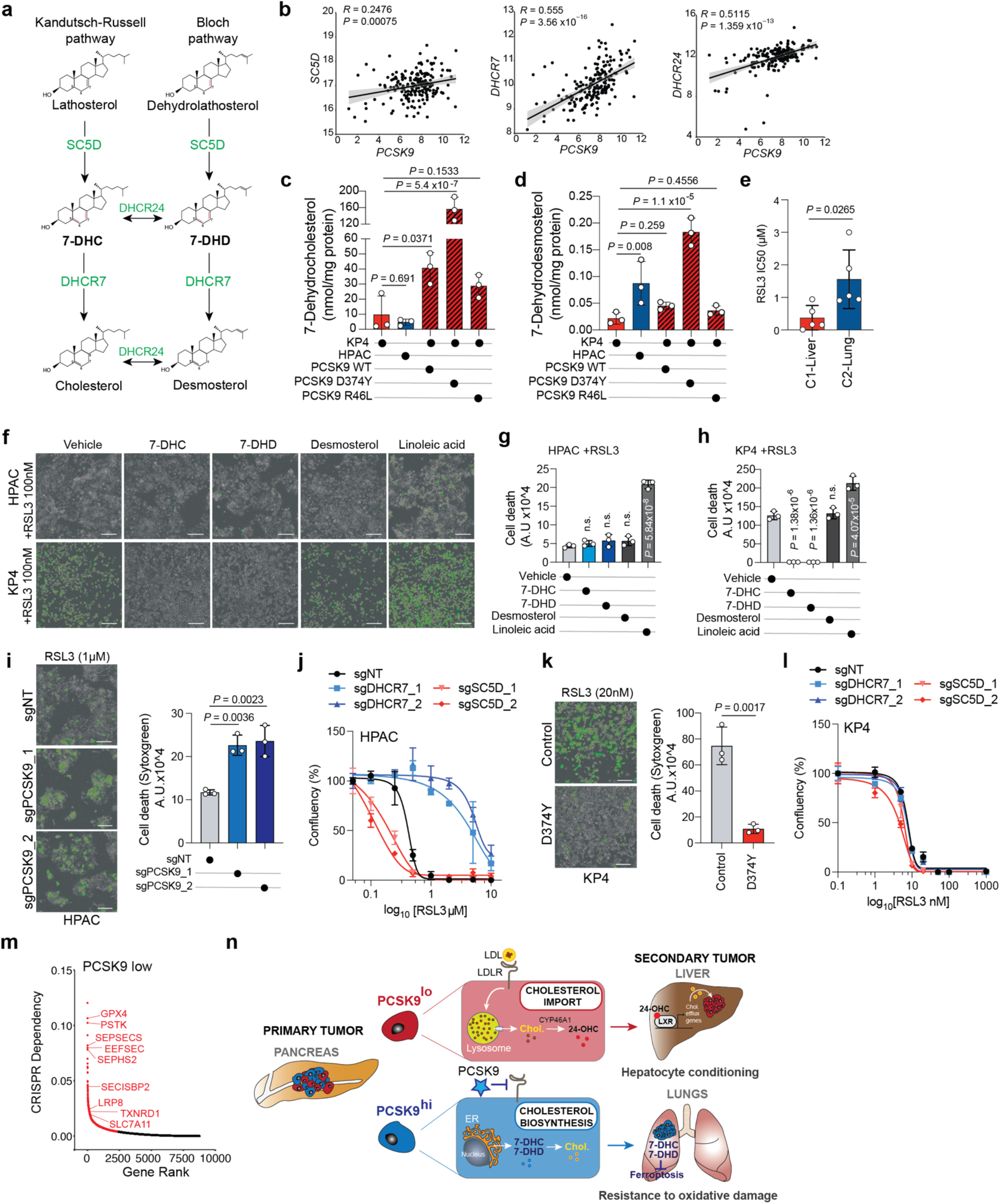
**a.** Schematic showing distal cholesterol biosynthesis steps that generate 7-dehydrocholesterol (7-DHC), 7-dehydrodesmosterol (7-DHD). Enzymes are shown in green. The 5,7 diene B ring structure is shown by the red lines. **b.** Scatter plot depicting correlation between high *PCSK9* transcript expression relative to distal cholesterol biosynthesis enzymes derived from TCGA PAAD datasets. **c, d.** Measurement of 7-DHC (c) and 7-DHD (d) in the indicated cell lines and conditions. N=3 biological replicates/condition. **e.** Average IC50 measurement for the GPX4 inhibitor, RSL3, for the C1-Liver (KP4, MiaPaCa-2, Panc-1, PSN1, PaTu8988T) and C2-Lung (HPAC, HPAF-II, CAPAN-2, YAPC and PaTu8902) lines. **f.** SYTOXgreen images depicting cell death (green fluorescence) of HPAC (top) and KP4 (bottom) cells treated with 100nM RSL3 plus vehicle or the indicated lipids for 24hrs. Scale bar: 200µm. **g.** Quantification of the SYTOXgreen images shown in F for HPAC. (n.s.; not significant). **h.** Quantification of the SYTOXgreen images shown in F for KP4. (n.s.; not significant). **i.** SYTOXgreen images (left) and quantification (right) of HPAC cell death (green fluorescence) following knockout of PCSK9 and treatment with RSL3 for 24hrs. Scale bar: 200µm. **j.** Measurement of HPAC cell growth following knockout of the indicated genes and treatment with increasing doses of RSL3 for 96hrs. **k.** SYTOXgreen images (left) and quantification (right) of KP4 cell death (green fluorescence) following expression of a control vector or PCSK9^D374Y^ and treatment with RSL3 for 24hrs. **l.** Measurement of KP4 cell growth following knockout of the indicated genes and treatment with increasing doses of RSL3 for 96hrs. **m.** DepMAP CRISPR dependency ranking of essential genes associated with PCSK9 low status across all cancers. (N = 1005 cell lines). **n.** Model depicting how the expression levels of PCSK9 dictate LDL-cholesterol import (top) versus biosynthesis (bottom), which in turn imparts metabolic fitness for growth in the liver or lungs, respectively.

To test whether differences in cholesterol biosynthesis pathway engagement is predictive of sensitivity to ferroptosis inducing agents in PDAC cells, we next measured the relative sensitivity of C1-Liver and C2-Lung lines to the Glutathione Peroxidase 4 (GPX4) inhibitors, RSL3 and ML210. C2-Lung lines were collectively more resistant to GPX4 inhibition, as measured by an overall higher mean IC50, relative to C1-Liver lines (**Fig. 5e; Extended Data Fig. 9b**). Similarly, HPAC cells were relatively resistant to RSL3 treatment (**Fig. 5,f,g**). In contrast, treatment of the C1-Liver line, KP4, with RSL3 caused a significant increase in cell death, as measured by SYTOXgreen (**Fig. 5f**). Importantly, sensitivity of KP4 cells to RSL3 was reversed following addition of exogenous 7-DHC or 7-DHD but not desmosterol (**Fig. 5f-h**), while addition of the polyunsaturated fatty acid (PUFA), linoleic acid, accelerated cell death in the presence of RSL3 in both KP4 and HPAC cells (**Fig. 5f-h**). These results indicate that reduced activity of the distal cholesterol biosynthesis pathway introduces a vulnerability to oxidative damage-inducing agents in C1-Liver cells. Accordingly, knockout of PCSK9 (**Fig. 5i**) or blocking the synthesis of 7-DHC and 7-DHD following knockout of SC5D in HPAC cells (**Fig. 5j**), led to a corresponding increase in sensitivity to GPX4 inhibitors, while suppression of DHCR7 and the resulting accumulation of 7-DHC and 7-DHD in HPAC cells, led to reduced sensitivity to GPX4 inhibition (**Fig. 5j**; **Extended Data Fig. 9c**). In contrast, ectopic expression of dominant active PCSK9^D374Y^ in KP4 cells, which causes a switch towards cholesterol biosynthesis (see **Figure 3j**), led to reduced sensitivity to GPX4 inhibition (**Fig. 5k; Extended Data Fig 9d**), while knockout of SC5D or DHCR7 had no effect (**Fig. 5l**). Consistent with these findings, unbiased analysis of CRISPR dependency data showed that PCSK9 low status across all cancer cell lines (N = 1005) is strongly associated with sensitivity to GPX4 loss, as well as several additional genes involved in protection against ferroptosis and lipid peroxidation (**Fig. 5m**). Taken together these data suggest that cells actively engaged in cholesterol biosynthesis have the added benefit of also producing anti-ferroptotic sterol intermediates, which enable more efficient colonization of a highly oxygenated organ such as the lung. On the other hand, cells which do not actively engage sterol biosynthesis are more vulnerable to oxidative damage induced cell death.

## Discussion

Most cancer cells exhibit increased synthesis, uptake and storage of lipids for growth and stress adaptation^40–43^. The optimal balance between these various lipid-handling activities may be partly dictated by energetic constraints: cholesterol synthesis is energetically expensive, thus cholesterol biosynthesis pathway genes are subjected to feedback regulation to maintain the levels of this lipid within a defined range compatible with growth^40,44^. Additionally, the mevalonate pathway and distal cholesterol synthesis pathways (referred to as the Bloch and Kandutsch-Russell pathways) produce several growth-promoting metabolites, including prenylation precursors, dolichol, ubiquinone^42^ and anti-ferroptotic derivatives^45^. Due to the high plasticity and regulation of the mevalonate-cholesterol pathway, the relative contributions of its branches and outputs to various aspects of cancer cell growth have remained unclear. Our study demonstrates that the differential utilization of the mevalonate-cholesterol pathway by PDAC cells can be best understood in the context of adaptation and colonization of metabolically diverse niches within the liver and lungs (**Fig. 5n**).

PCSK9 emerges as a key determinant of secondary organ colonization preference in PDAC. While extensive studies of PCSK9, conducted in the context of cardiovascular disease, have uncovered its role as a master regulator of LDLR stability and LDL-cholesterol import, far less is understood about its roles in the altered metabolism of cancer cells^46^. We demonstrate that the ability of PSCK9 to switch PDAC cell metabolism between cholesterol import (PCSK9 low status) versus synthesis (PCSK9 high status), is sufficient to impart different colonization fates. A key finding from our studies is that PCSK9 levels do not simply dictate the rate of uptake versus de novo synthesis of cholesterol. Instead, PCSK9 levels impart completely distinct patterns of utilization of mevalonate-cholesterol intermediates and derivatives, whereby liver-avid, PCSK9-low cells convert cholesterol into 24-OHC to reprogram the microenvironment and induce nutrient release from neighboring hepatocytes, whereas PCSK9-high PDAC cells utilize 7-DHD and 7-DHC, intermediates with powerful free radical trapping capabilities, to prevent ferroptosis in the oxygen rich environment of the lung. This finding explains why liver-avid cells grow less efficiently in the lung – likely due to their inability to produce 7-DHD and 7-DHC. Accordingly, increasing PCSK9 levels in liver-avid cells, which leads to a corresponding switch towards cholesterol biosynthesis, enables their growth in the lung. Likewise, PCSK9 ablation in lung-avid cells causes a shift towards cholesterol import and enables grow in the liver. These findings establish PCSK9 as necessary and sufficient for secondary organ site preference and highlight how differential modes of cholesterol acquisition and utilization can enable adaptation to growth in metabolically diverse metastatic niches (**Fig. 5n**).

A relationship between cholesterol metabolism and PDAC differentiation has been proposed based on prior transcriptional and function studies^47–49^. For instance, high expression levels of cholesterogenic genes in PDAC cells correlates with the classical subtype and overall better prognosis^47,49^, which we also observe in our study. On the other hand, high levels of LDLR was associated with aggressive disease and an increased risk of PDAC recurrence^50^. A switch from classical to basal differentiation was also shown to occur following deletion of *NSDHL* – a distal cholesterol biosynthesis gene required for generation of zymosterone – in a mouse model of PDAC^48^. Our studies suggest that PCSK9 is an important node that links cholesterol status with differentiation and the related subtype specific behaviors.

Several large-scale studies evaluating the potential benefit of cholesterol lowering drugs (eg. Statins) on cancer incidence and mortality have reported different findings based on population distribution, cancer type, duration of inhibitor use or pre-existing cardiovascular disease^41,46,51,52^. Moreover, conflicting reports on the benefits of statin use as therapy for advanced cancer have made clinical translation of cholesterol-lowering agents challenging. However, a recent study evaluating advanced ovarian, lung and pancreatic cancer showed a significant correlation between statin use and decreased risk of mortality^53^. Whether a demonstrable link between PCSK9 inhibitor use and incidence of liver metastases and patient outcomes exists remains to be determined.

Finally, our study suggests that a more nuanced manipulation of the cholesterol biosynthetic pathway could achieve greater ability to exploit specific vulnerabilities of cancer cell populations and patients. For instance, interventions that block tumor cell-hepatocyte cross talk by targeting 24-OHC production or LXR activation could antagonize liver colonization. Likewise, the inability of liver-avid PDAC cells to generate 7-DHC and 7-DHD make them more vulnerable to ferroptosis inducing agents, a finding with potential clinical applications. Conversely, the elevated dependency of lung-avid PDAC cells on 7-DHC and 7-DHD suggest that directly blocking their production may antagonize lung colonization.

## Extended data figure legends

**Extended Data Figure 1.**
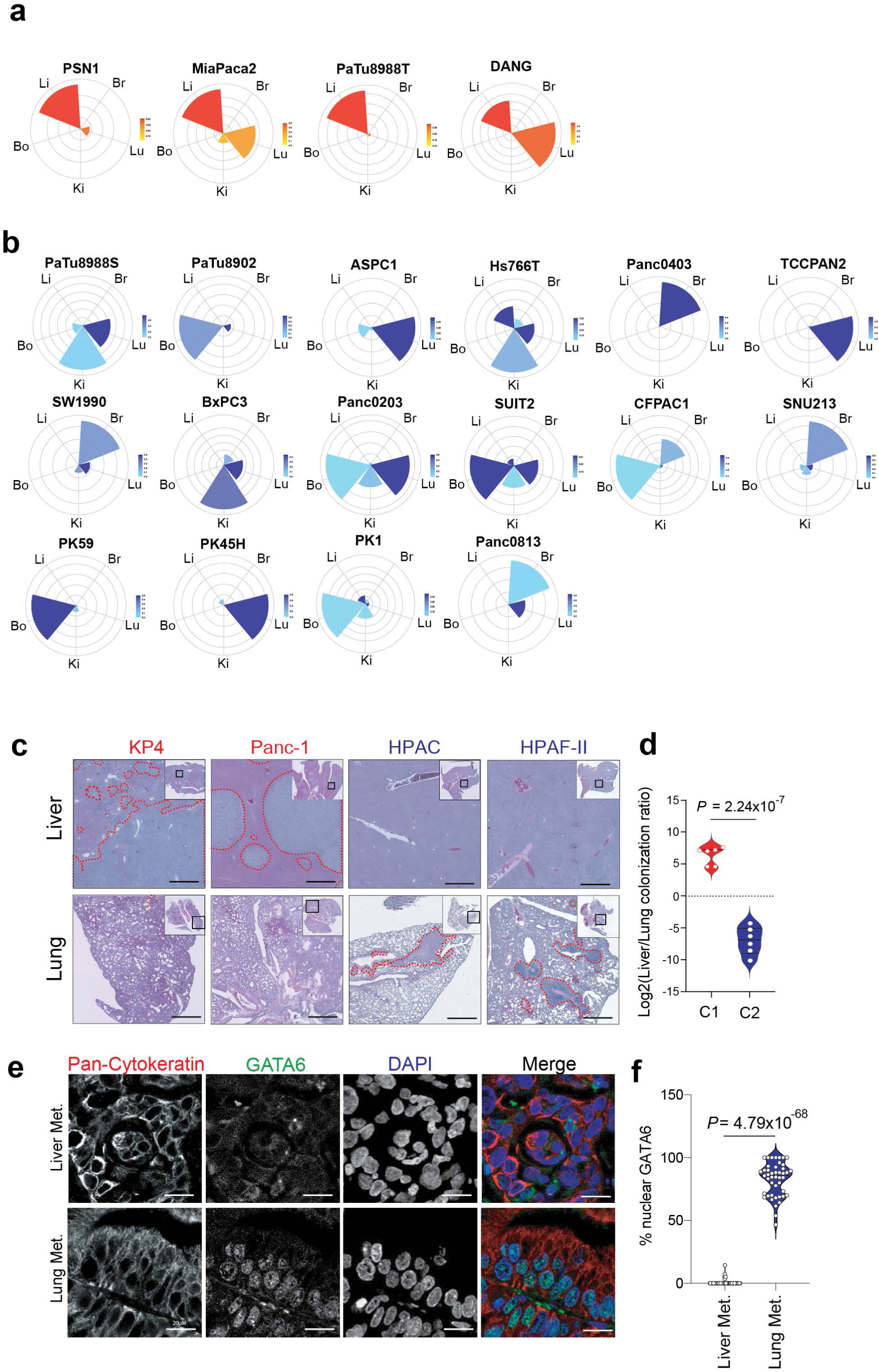
**a, b.** Petal plots depicting metastatic potential (length) and penetrance (width) for the C1-Liver lines (a) and C2-Lung lines (b), extracted from the MetMap 500 analysis. Br, Brain; Bo, Bone; Ki, Kidney; Li, Liver; Lu, Lung. **c.** H&E staining showing tumor growth (red outline) in the liver (top) and lung (bottom) following intracardiac injection of the indicated PDAC cell lines. Scale bars: 500 µm. **d.** Quantification of liver/lung colonization ratio of C1-Liver and C2-lung lines for the experiment in c. N = 6 mice per/cluster. **e.** Immunofluorescence images of GATA6 (green), Pan-Cytokeratin (CK19) (red), DAPI (blue) in liver and lung patient PDAC metastatic lesions. Note the nuclear localization of GATA6 in the lung metastases. Scale bars: 20 µm. **f.** Quantification of the percentage nuclear GATA6 in liver and lung metastases sections (N=30 total fields from 3 liver and 3 lung metastases sections).

**Extended Data Figure 2.**
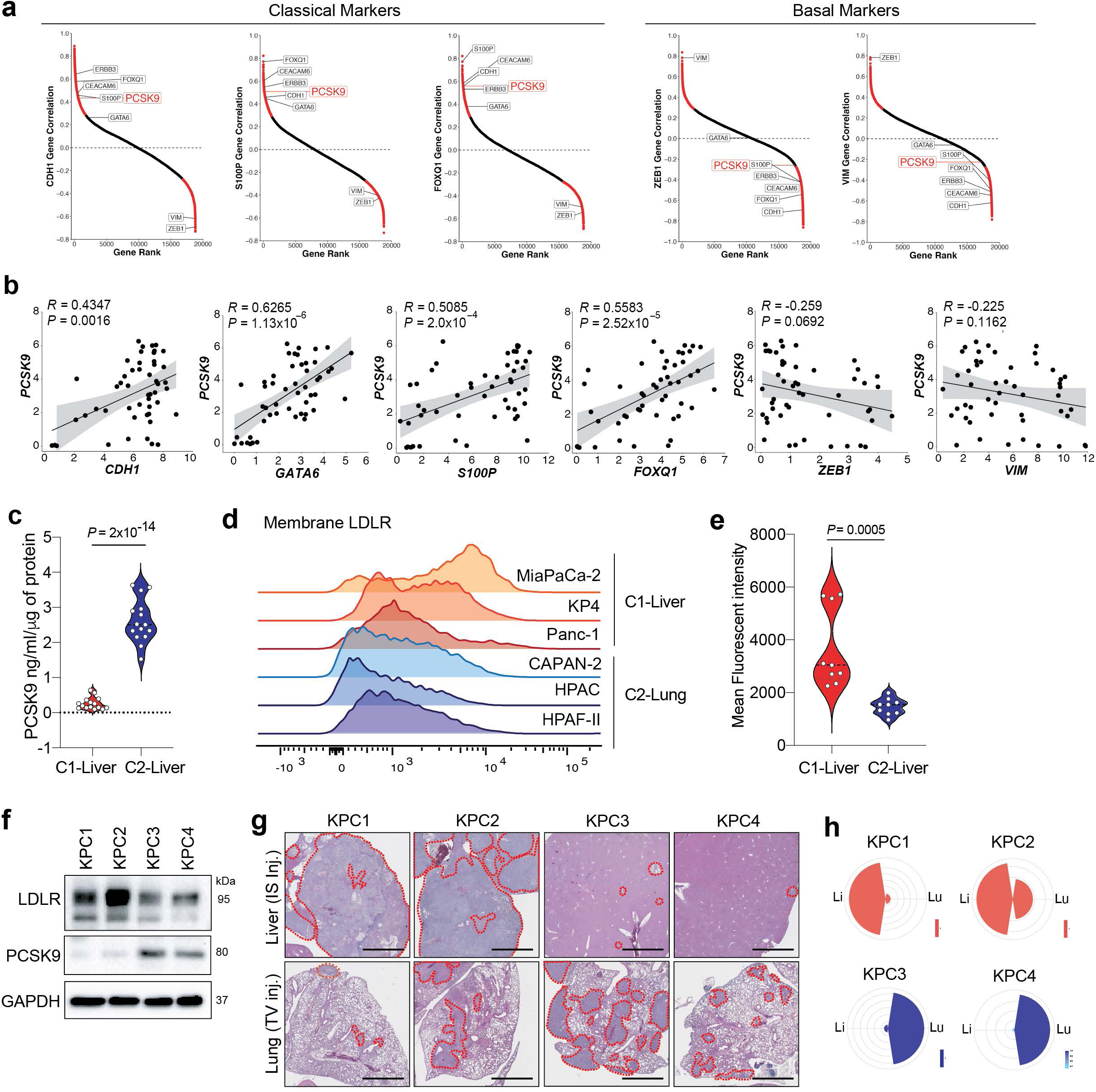
**a.** Gene expression correlation between PCSK9 the indicated classical (*S100P, FOXQ1, CDH1*) and basal (*VIM, ZEB1*) genes across all PDAC cells lines available on DepMAP. Significant correlations are indicated in red, and non-significant correlations are indicated in black. *PCSK9* and classical (*CDH1, ERBB3, FOXQ1, CEACAM6, S100P, GATA6*) and basal (*ZEB1, VIM*) genes are indicated for each plot. **b.** Scatter plot showing expression correlation between PCSK9 and individual classical (*CDH1, GATA6, S100P, FOXQ1*) and basal (*ZEB1, VIM*) genes expression (data extracted from n=50 cell lines available in DepMap). **c.** ELISA based quantification of secreted PCSK9 protein from conditioned media collected from C1-Liver (KP4, MiaPaCa-2, Panc-1, PSN1, PaTu8988T) and C2-Lung (HPAC, HPAF-II, CAPAN-2, YAPC and PaTu8902) cell lines. (N=3 biological replicates). **d.** Flow cytometry-based detection of plasma membrane LDLR in three C1-Liver (MiaPaCa-2, KP4 and Panc-1) and three C2-Lung (CAPAN-2, HPAC and HPAF-II) cell lines. **e.** Quantification of the mean fluorescent intensity of plasma membrane LDLR from the cell lines in d (N=3 biological replicates per cell line). **f.** Western blot of the indicated proteins in mouse derived KPC cell lines. **g.** H&E staining of liver (top) and lung (bottom) following intrasplenic or tail vein injection, respectively, of the indicated KPC cell lines. N=3 mice/conditions. Scale Bar: 500µm **h.** Petal plots depicting metastatic potential (length) and penetrance (colour) for the indicated cell lines. Li, Liver; Lu, Lung.

**Extended Data Figure 3.**
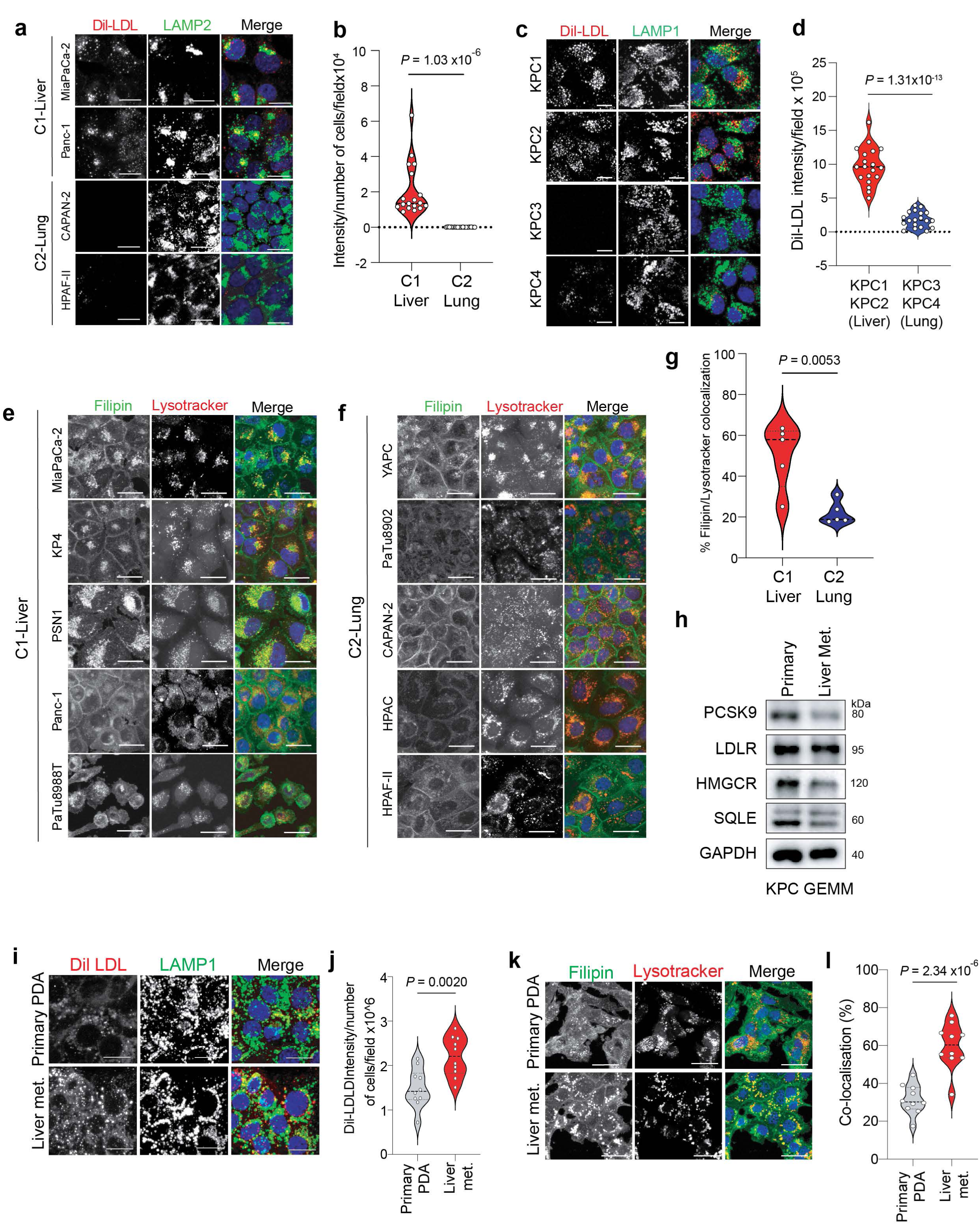
**a.** Immunofluorescence images depicting Dil-LDL uptake (red) in C1-Liver (MiaPaCa-2, Panc-1) and C2-Lung (CAPAN-2, HPAF-II) lines. Co-staining with LAMP2 (green) highlights lysosomal localization of the internalized lipoprotein (yellow). Scale bars: 10 µm (N=10 fields per cell line). **b.** Quantification of staining intensity per cell per field of data depicted in a. **c.** Immunofluorescence images depicting Dil-LDL uptake (red) and co-staining for LAMP1 (green) in four mouse KPC derived cell lines. Note that KPC1 and KPC2 take up exogenous LDL while KPC3 and KPC4 do not. Scale bars: 10 µm (n=10 fields per cell line). **d.** Quantification of staining intensity per cell per field of data depicted in c. **e,f.** Immunofluorescence images depicting Filipin (green) in C1-Liver (MiaPaCa-2, KP4, PSN1, Panc-1, PaTu8988T) (e) and C2-Lung (YAPC, PaTu8902, CAPAN-2, HPAC, HPAF-II) (f) lines. Scale bars: 10µm. **g.** Quantification of filipin/lysotracker co-localization per cell per field for data depicted in e and f. **h.** Western blot for the indicated proteins in KPC derived isogenic primary and liver metastases cells. **i.** Immunofluorescence images depicting Dil-LDL uptake (red) and co-staining for LAMP1 (green) in the KPC lines in h. Scale bars: 10µm. **j.** Quantification of staining intensity per cell per field of data depicted in i. (N=10 fields/cell line). **k.** Immunofluorescence images depicting Filipin (green) and Lysotraker (red) in the KPC lines in h and I. Scale bars: 10µm. **l.** Quantification of filipin/lysotracker co-localization per cell per field for data depicted in k. (n=10 fields/cell line)

**Extended Data Figure 4.**
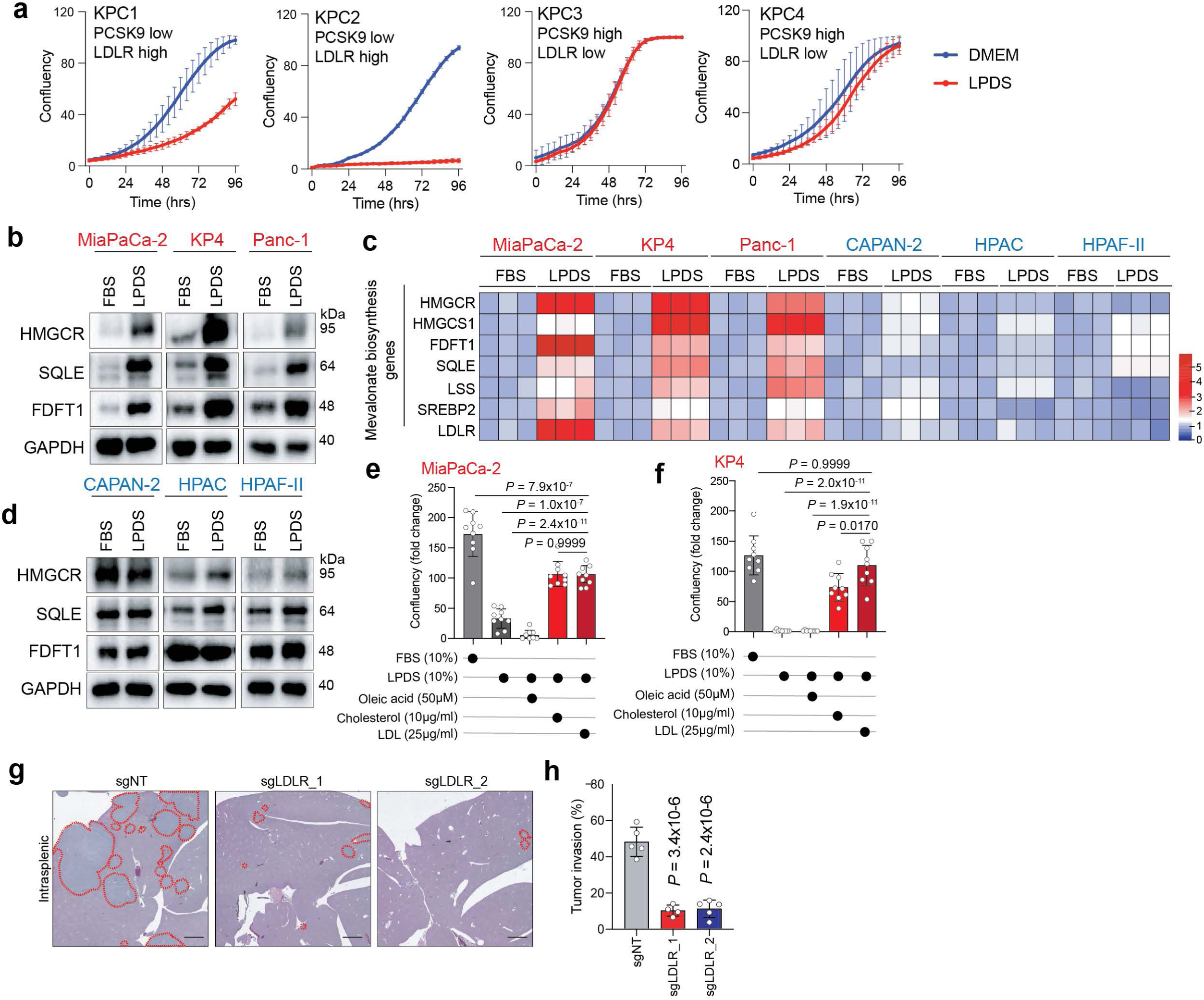
**a.** Growth curves of KPC lines cultured in DMEM and standard FBS or DMEM containing lipoprotein depleted serum (LPDS). **b.** Western blot of indicated proteins in C1-Liver (MiaPaca-2, KP4, Panc1) following treatment with DMEM and standard FBS or DMEM containing LPDS for 72 hrs. **c.** Heatmap depicting the expression levels of mevalonate pathway genes in C1-Liver (MiaPaca-2, KP4, Panc1) and C2-Lung (CAPAN-2, HPAC, HPAF-II) lines. Each row represents a specific gene and 3 biological replicates are shown for each cell line. Expression levels are expressed as zscores. **d.** Western blot of indicated proteins in C2-Lung (CAPAN-2, HPAC, HPAF-II) lines following treatment with DMEM and standard FBS or DMEM containing LPDS for 72 hrs. **e,f.** Relative cell confluency of MiaPaCa-2 (e) and KP4 (f) cells cultured in the indicated media showing supplemented with cholesterol, LDL or oleic acid. **g.** H&E staining showing tumor growth (red outline) in the liver 28 days post intra-splenic injection of KP4 cells transfected with control sgRNA or sgRNA targeting LDLR. Scale bars: 200 µm. **h.** Quantification of tumor area from data in g. N=5 mice/condition.

**Extended Data Figure 5.**
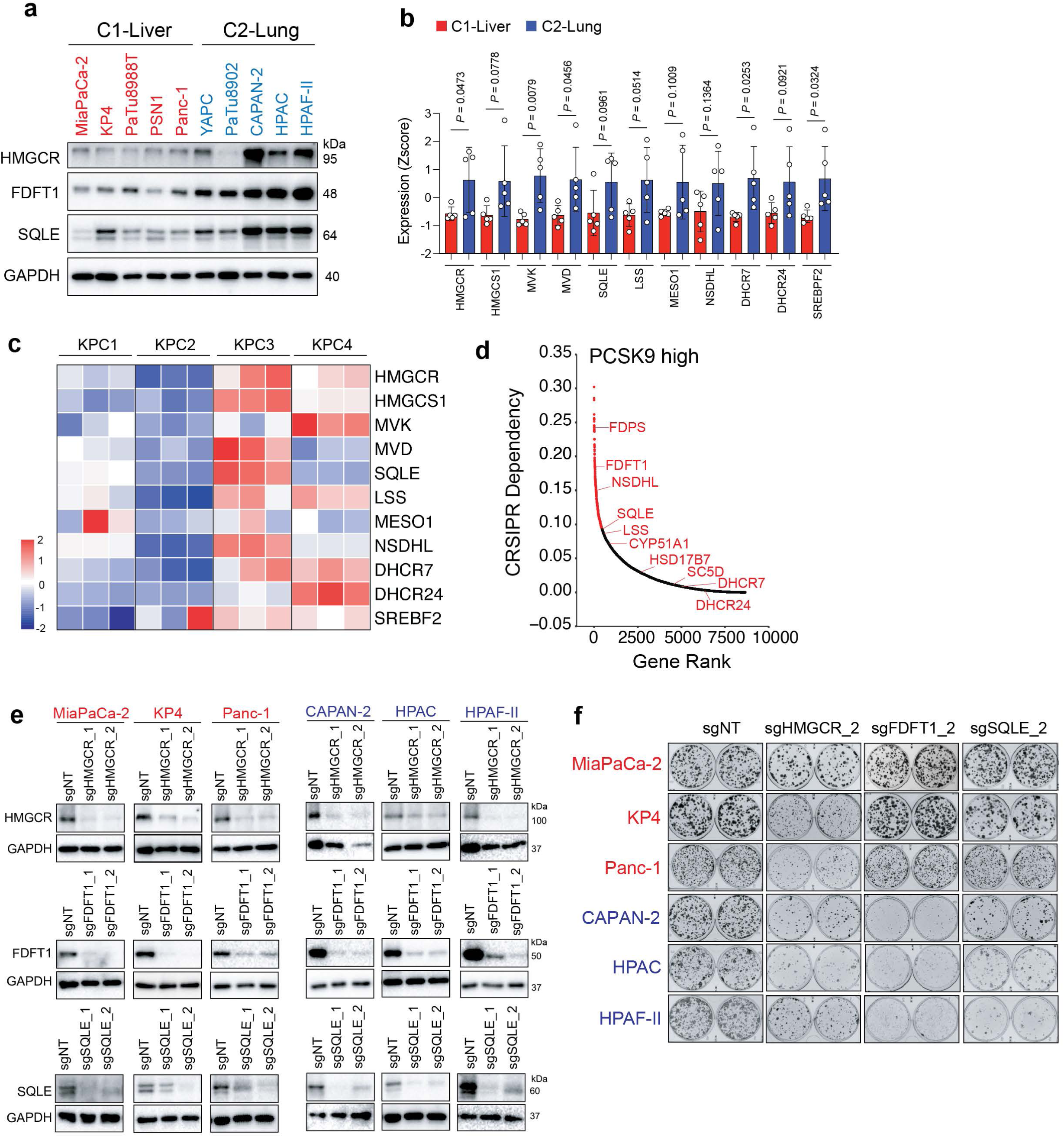
**a.** Western blot of the indicated proteins in C1-Liver and C2-Lung lines. **b.** Quantitative real-time PCR measurement of the indicated genes in C1-Liver (N=5) and C2-Lung (N=5) lines expressed as z-score. Each point represents the average of N=3 biological replicates for each line. **c.** Heatmap depicting mRNA expression of the indicated genes displayed as z-score. N=3 biological replicates per line. **d.** DepMAP CRISPR dependency ranking of essential genes associated with *PCSK9* high status across PDAC. **e.** Western blot for the indicated proteins following CRISPR mediated knockout of the indicated genes (related to data presented in figure 3f,h). **f.** Colony formation in C1-Liver (MiaPaCa-2, KP4, Panc-1) and C2-Lung (CAPAN-2, HPAC, and HPAF-II) lines upon CRISPR-mediated knockout using a second independent sgRNA targeting the indicated genes. Note that the sgNT images are identical to that depicted in figure 3f,h.

**Extended Data Figure 6.**
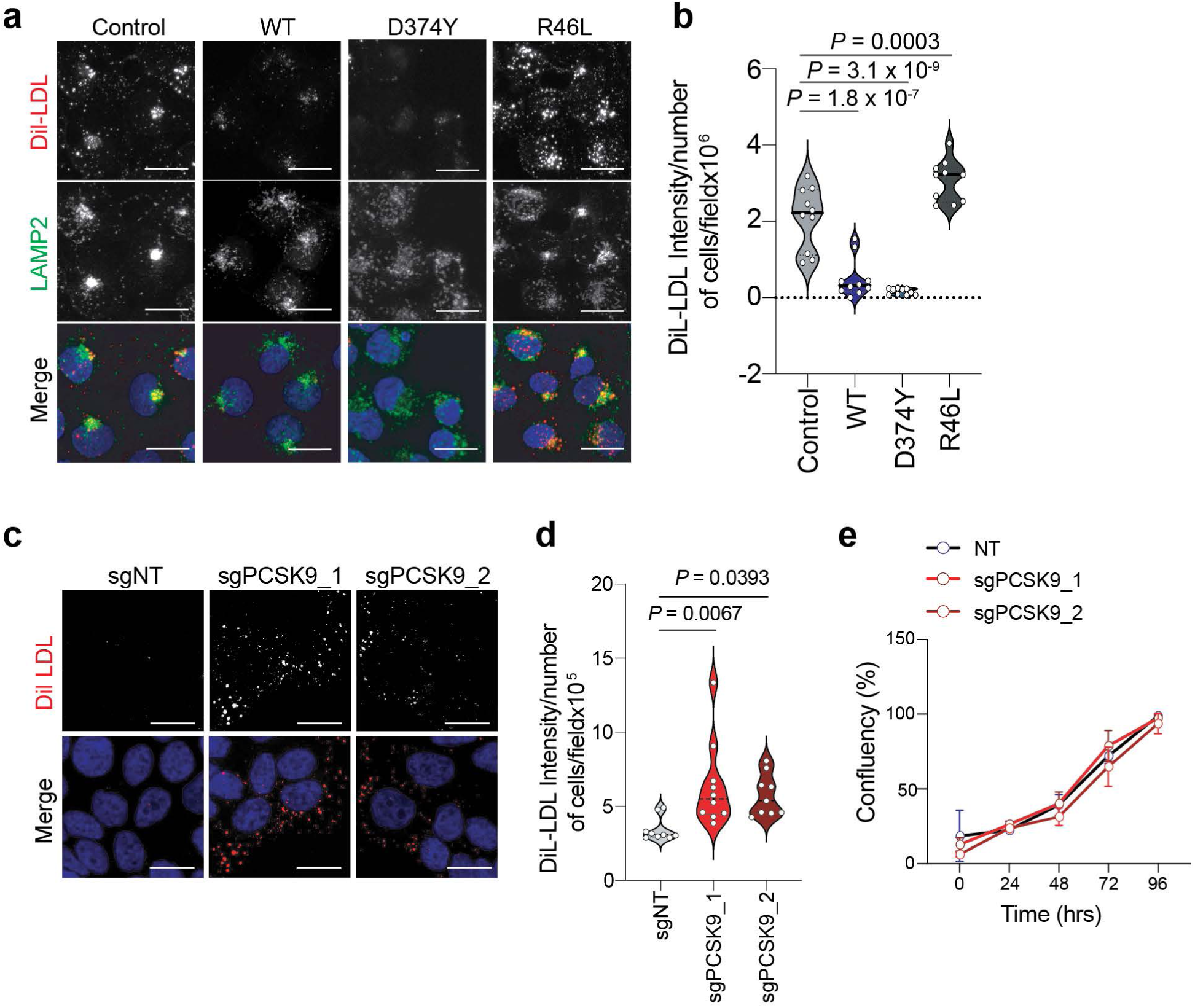
**a.** Immunofluorescence images of DiL-LDL (red) uptake and LAMP2 (green) in KP4 parental cells or following ectopic expression of the indicated PCSK9 variants. Scale bar: 10µm. **b.** Quantification of DiL-LDL fluorescence intensity per cell per field from each condition depicted in A. N=10 fields/cell lines. **c.** Immunofluorescence images of DiL-LDL (red) uptake in control HPAC cells (sgNT) or following knockout of PCSK9 (sgPCSK9). Scale bar: 10µm. **d.** Quantification of DiL-LDL fluorescence intensity per cell per field from each condition depicted in c. N=10 fields/cell lines. **e.** Measurement of percentage confluency of HPAC cells transfected with sgNT or sgPCSK9.

**Extended Data Figure 7.**
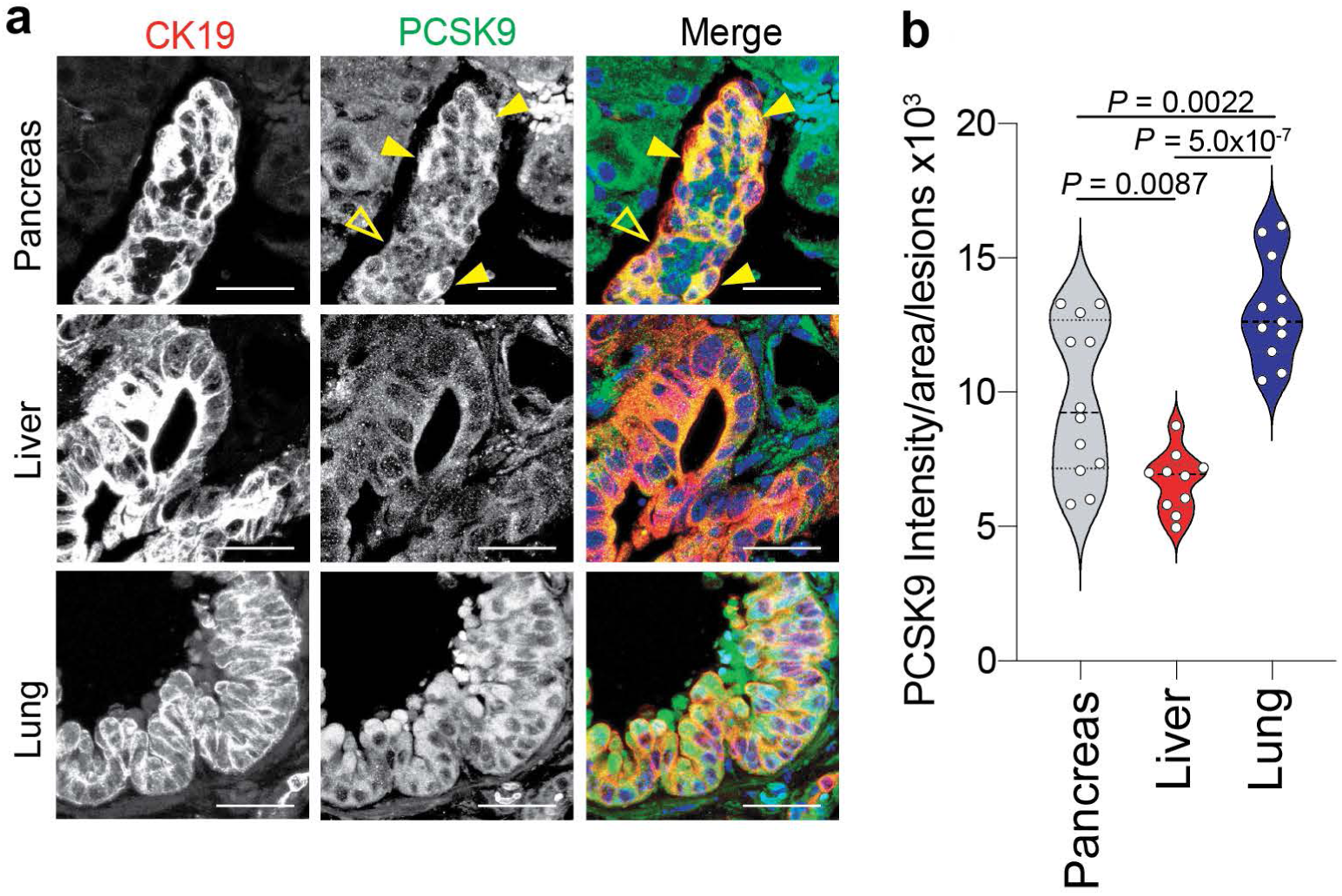
**a.** Immunofluorescence images depicting CK19 (red) and PCSK9 (green) in mouse MT23 cells injected orthotopically into the pancreas, via intra-splenic injection to seed the liver and via tail-vein injection to seed the lungs of C57Bl/6 mice. Scale bars: 50 µm. **b.** Quantification of PCSK9 staining intensity in CK19 positive ductal structures from tumours grown in the pancreas (N=3 mice; 12 fields), liver (N=3 mice; 10 fields) and Lung (N=3 mice; 12 fields).

**Extended Data Figure 8.**
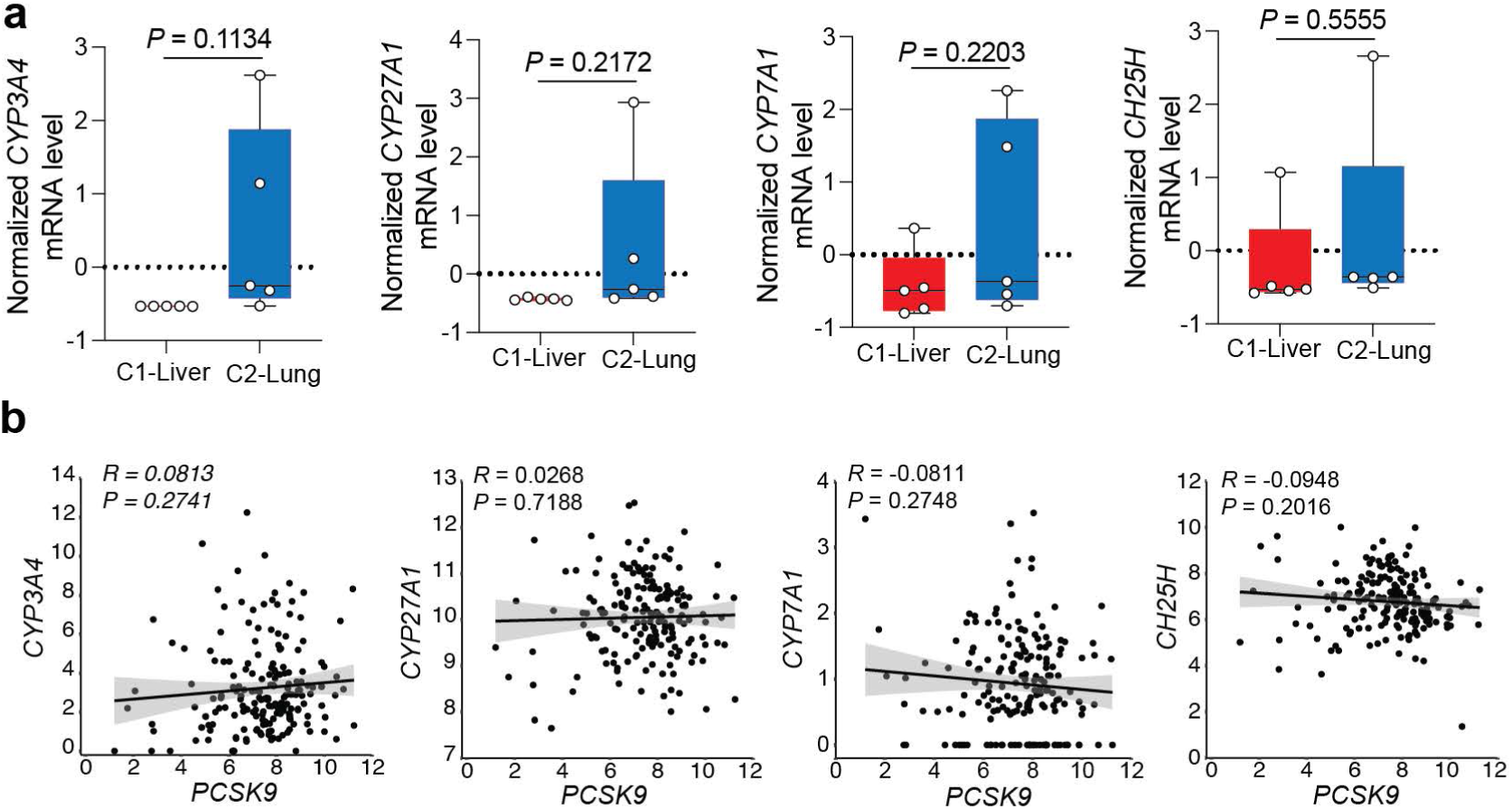
**a.** Normalized mRNA expression measured by quantitative real-time PCR of the indicated transcripts in C1-Liver (KP4, MiaPaCa-2, Panc-1, PSN1, PaTu8988T) and C2-Lung (HPAC, HPAFII, CAPAN-2, YAPC and PaTu8902) lines. Dots represent the average of 3 biological replicates per line. **b.** Scatter plots showing TCGA PAAD derived expression correlation between *PCSK9* and the indicated genes.

**Extended Data Figure 9.**
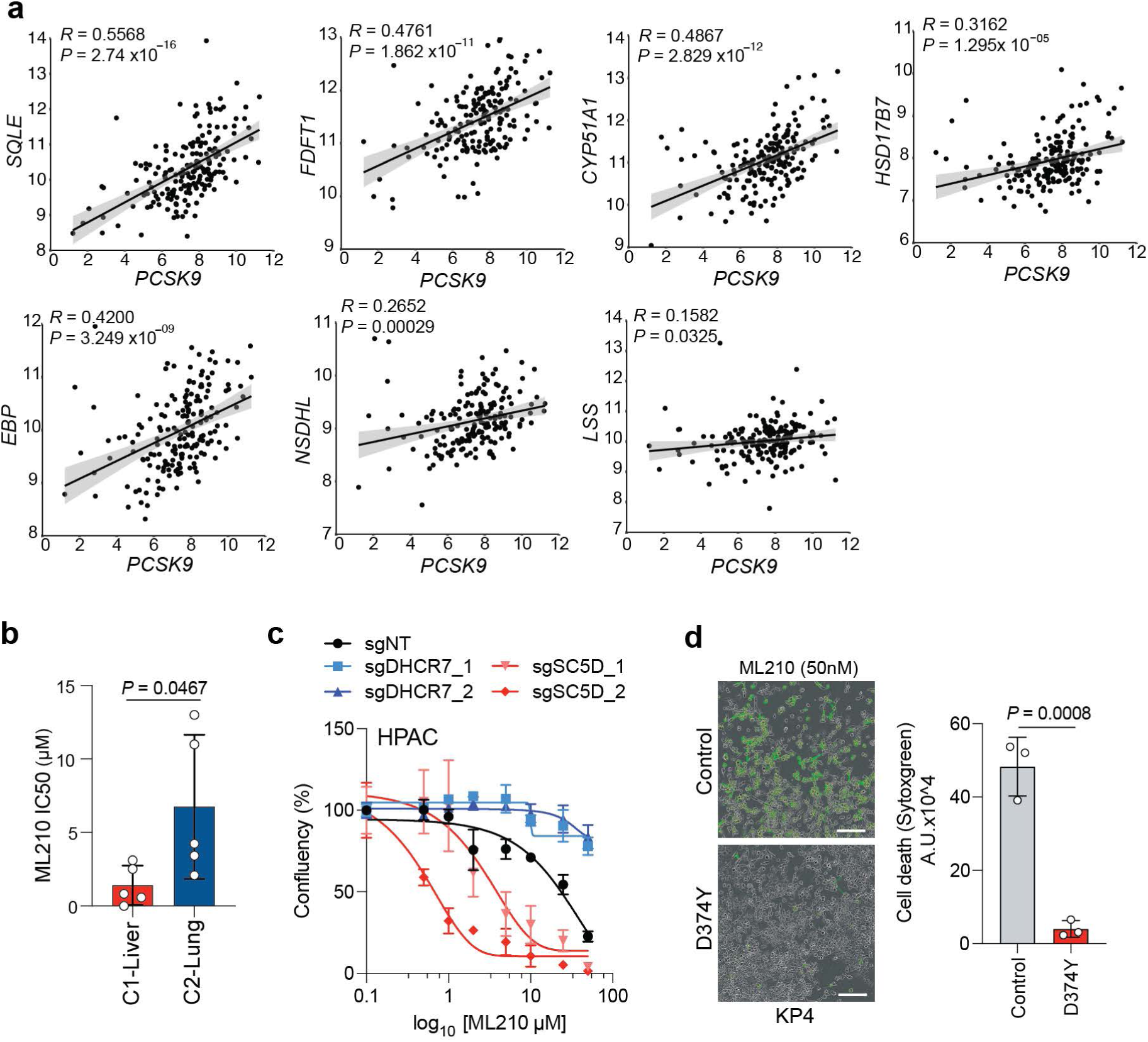
**a.** Scatter plots showing TCGA PAAD derived expression correlation between PCSK9 and individual cholesterol biosynthesis pathway genes. **b.** Average IC50 measurement for the GPX4 inhibitor, ML210, in C1-Liver (KP4, MiaPaCa-2, Panc1, PSN1, PaTu8988T) and C2-Lung (HPAC, HPAF-II, CAPAN-2, YAPC and PaTu8902) lines. **c.** Measurement of HPAC cell growth following knockout of the indicated genes and treatment with increasing doses of ML210 for 96hrs. **d.** SYTOXgreen images and quantification of KP4 cell death (green fluorescence) following expression of a control vector or PCSK9^D374Y^ and treatment with ML210 for 24hrs.

## Materials and Methods

### Cell Culture and Reagents

MiaPaCa-2, KP4, PaTu8988T, PSN1, Panc-1, YAPC, PaTu8902, CAPAN-2, HPAC, HPAF-II and HEK-293T were obtained from the American Type Culture Collection (ATCC) or DSMZ. MT23 and MM10 cells were established from Pdx1-cre+; KrasLSL-G12D/+; Trp53lR172H/+ mice, backcrossed into a C57BL/6 background, and isolated from the pancreas and liver, respectively (provided by Dr. Chang-II Hwang, University of California, Davis). FC1245 (KPC1) were provided by Dr. David Tuveson (CSHL); HY15549 (KPC2) and HY19636 (KPC3) were provided by Dr. Ronald DePinho (MD Anderson); MT3 (KPC4) were provided by Dr. Chang-il Hwang and Dr. David Tuveson and were all derived from Pdx1-cre+; KrasLSL-G12D/+; Trp53 R172H/+ mice. All cell lines were cultured in DMEM media (Gibco) supplemented with 10% FBS (Atlanta biologicals), 1% Pencillin/Streptomycin (Gibco) and 15mM HEPES (Gibco) and grown in a humidified chamber at 37°C, 5% CO_2_. Cells were trypsinized using TrypLE (Gibco). Routine mycoplasma testing was performed using the Mycoplasma PCR detection kit (Abm; Cat. Number G238) at least once a month and the cell lines were authenticated by STR fingerprinting. Cell lines are passaged for a maximum of 15 passages upon thawing prior to replacement. Dil-LDL was purchased from Thermofisher (Cat. Number L3482), Filipin was purchased from Sigma (Cat. Number F976550MG), Lysotracker was purchased from ThermoFischer (Cat. Number L7528). Lipoprotein depleted serum was purchased from Kalen Biomed (Cat. Number 880100-1). 24hydroxycholesterol was purchased from Sigma (Cat. Number SML-1648). SYTOXgreen was purchased from Thermofisher scientific (Cat. Number S7020). 7-dehydrodesmosterol (Cat. Number EVU144) and desmosterol (Cat. Number EVU126) were purchased from Kerafast Inc, 7dehydrocholesterol (Cat. Number 30800-5G-F) and Linoleic acid (Cat. Number L1012-1G) were purchased from Sigma.

#### Plasmids

pLX304 Luciferase-V5 blast was a gift from Kevin Janes (Addgene plasmid # 98580) and pAAV/D374Y-hPCSK9 was a gift from Jacob Bentzon (Addgene plasmid # 58379). To generate the wild-type and R46L PCSK9 variants, the full-length sequence was first cloned into pJML1 and Quickchange mutagenesis (Agilent Cat. Number 210519-5) was used to introduce the point mutations followed by sequence variation.

### Lentiviral experiments

Lentivirus was produced by transfecting HEK293T cells with lentiviral vector pLKO.1 containing shRNA sequence and packaging plasmids psPAX2 (Addgene, plasmid #: 12260) and pMD2.G (Addgene, plasmid #: 12259) at a ratio of 0.5:0.25 using X-tremeGENE transfection (Sigma Aldrich, Cat. Number 6365787001) reagent following manufacturer’s instructions. The supernatant containing the virus was collected after 48h by passing through 0.45µM filter to remove cellular debris. Target cells were infected with the virus-containing media using Polybrene reagent (EMD Millipore, Cat. Number TR-1003-G) following the manufacturer’s protocol and selected for 48 hours in 2 µg/ml puromycin.

### Immunoblotting

Cells were lysed in ice-cold buffer (50 mM HEPES, pH 7.4, 40 mM Sodium Chloride, 10 mM Sodium-pyrophosphate, 10 mM ß-glycerophosphate, 50 mM Sodium fluoride, 2 mM EDTA, and 1% Triton X-100, supplemented with protease inhibitor tablets (Fisher Scientific, Cat. Number A32965). Protein concentration was determined using a Pierce BCA protein assay kit (Life Technologies, Cat. Number 23227), and 20 µg of protein was resolved on 12% acrylamide gels and subsequently transferred onto PVDF membranes (EMD Millipore, Cat. Number IPVH00010). Following blocking with 5% skim milk in Tris-buffered saline with 0.2% Tween (TBS-T) (blocking buffer), membranes were incubated overnight at 4°C with primary antibodies in blocking buffer. The membranes were then washed three times with TBS-T and further incubated with species-specific horseradish peroxidase-conjugated secondary antibodies for one hour in blocking buffer. Membranes were developed using SuperSignal West Pico chemiluminescent substrate (Fisher Scientific, Cat. Number 34080) and imaged with the ChemiDoc XRS+ system (Bio-Rad). Antibodies used are listed in Supplementary Table 7.

### Proliferation assays

For clonogenicity assays, KP4 and MiaPaca-2 cells were seeded at 2000 cells per well, Panc-1 cells at 4000 cells per well, and CAPAN, HPAC, and HPAF cells at 8000 cells per well in 6 well plates in growth media. After 10 days, the growth medium was aspirated, and cells were washed with PBS, fixed with ice-cold methanol for 10 mins. and stained with crystal violet solution (0.1% w/v) for 15 mins at room temperature. For experiments using lipoprotein-depleted serum (LPDS), cells were seeded in 96-well plates cells per well in growth media and allowed to grow for 24hrs (KP4 and MiaPaCa-2 at 2000 cells per well, Panc-1 at 4000 cells per well, CAPAN-2, HPAC, and HPAF cells at 8000 cells per well). Subsequently, the cells were washed with PBS and incubated in fresh growth media or DMEM supplemented with LPDS for 96 hrs. At the conclusion of the assay, the medium was aspirated, and cells were washed with PBS and fixed with ice cold methanol for 10 minutes and stained with crystal violet solution (0.1% w/v). For proliferation assay with RSL3 treatment, cells were seeded in 96 well plates (4,000 cells per well) and treated with different concentration of RSL3 for 96 hours. Cells were then washed with PBS, fixed in ice cold methanol, and stained with crystal violet solution (0.1%w/w).

### SYTOXgreen assay

Cells were seeded at a concentration of 4,000 cells per well in a 96-well plate and treated with the different lipid metabolites for 24 hours at a concentration of 50µM. RSL3/ML210 treatment was then added and cell death was assessed using SYTOXgreen (dilution 1/30 000) for the indicated time. Green staining area was assessed and quantify by Incucyte S3.

### Immunohistochemistry

Slides were baked at 60°C for 1 hr, and paraffin removal was achieved through two 5-minute washes in xylene, followed by sequential rehydration in ethanol (100, 90, 70, 50, and 30%) for 5 mins each. After two water washes, samples were heat-treated for 20 mins in a 10 mM sodium citrate buffer solution (pH 6.0), washed twice with PBS, and incubated for 30 mins in methanol/5% hydrogen peroxide at room temperature to block endogenous peroxidase activity. Following two PBS washes, samples were blocked with 2.5% normal goat serum (NGS) for 1 hr and then incubated overnight with primary antibody (Supplementary Table 7) at 4°C. Subsequently, two 5 min PBS washes preceded incubation with secondary antibody. After three washes, slides were stained with di-aminebenzidine (DAB) substrate kit (Vector Laboratories, Cat. Number SK-4100) for 10 mins, washed in water, and counterstained with hematoxylin before dehydration and mounting. Bright field images were captured using a KEYENCE BZ-X710 microscope. Tumour sections were evaluated for staining positivity by a pathologist (KWW) and quantified using a modified histoscore ranging from 1 to 5 (1 = absent, 2 = weak, 3 = moderate, 4 = strong, and 5 = very strong).

### Immunofluorescence

Human and mouse cells were cultured on fibronectin-coated coverslips for 2 days after which cells were fixed with 4% PFA for 15 mins at room temperature, permeabilized with 0.1% saponin, and blocked with Normal Goat Serum (NGS) for 15 mins. Subsequently, samples were incubated overnight at 4°C with primary antibodies (Supplementary Table 7), then washed three times with PBS and incubated with secondary antibodies (Supplementary Table 7) at room temperature for 1 hour. Coverslips were then mounted on glass slides using DAPI Fluoromount-G (Southern Biotech, Cat. Number 0100-20), except for Filipin experiments where cells were incubated with Sytox green (ThermoFischer Cat. Number S7020) for 10 mins, washed three times, and directly mounted with Fluoromount-G (Southern Biotech, Cat. Number 0100-01). For Dil-LDL uptake assays, cells were cultured in serum-free medium for 16 hrs prior to incubation on ice for 30min with Dil-LDL (ThermoFisher Scientific, Cat. Number. L3482). The medium was then replaced with warmed fresh growth medium and cells were transferred to a 37°C incubator for varying durations. At the conclusion of the assay, coverslips were fixed and permeabilized as above and counter stained for LAMP2 and DAPI. For visualization of cholesterol, cells were first incubated with Lysotracker Red DND-99 (ThermoFisher Scientific, Cat. Number L7528) at a concentration of 100 nM for 30 mins at 37°C. Cells were then fixed with 4% PFA, permeabilized with 0.1% saponin and subsequently incubated with Filipin (200 µg/ml) for 30 mins at room temperature. After incubation, cells were washed with PBS and incubated with Sytox green (ThermoFischer Cat. Number S7020) at a dilution of 1:10,000 to label specific nuclei and mounted using Fluoromount-G mounting medium (Southern Biotech, Cat. Number 0100-01)

Immunofluorescence of tissue sections was undertaken by first baking the samples at 60°C for 1 hr, followed by paraffin removal through two 5 min xylene washes and sequential rehydration in ethanol (100, 90, 70, 50, and 30%) for 5 mins each. After two water washes, samples were heattreated for 20 mins in a 10 mM sodium citrate buffer solution (pH 6.0), washed twice with PBS, and then incubated for 30 mins in methanol at room temperature, followed by two PBS washes. Blocking was performed with 2.5% NGS for 1 hr after which samples were incubated overnight with primary antibody at 4°C (Supplementary Table 7). After two 5 min PBS washes, samples were incubated with the secondary antibody and mounted with DAPI Fluoromount-G. Stained cells and tissue sections were visualized using a Zeiss LSM900 Axio Observer Z1/7.

### RNA Extraction and quantitative PCR

Total RNA was isolated using PureLink RNA mini kit (Thermo Fisher, Cat. Number 12183025). cDNA synthesis was performed using the iScript reverse transcription supermix (Bio-Rad, Cat. Number 1708841), followed by real time quantitative PCR using iTaq universal SYBR Green Supermix (Bio-Rad, Cat. Number 1725122) on a CFX384 Touch Real Rime PCR Detection System (Bio-Rad). Primers used are detailed in Supplementary Table 8.

### CRISPR Knock-Out

CRISPR knockouts were generated employing the ribonucleoprotein-electroporation method as previously outlined by Liang et al. (2015) [cite: Liang J biothechnol 2015]. In brief, 200,000 cells were electroporated using an AMAXA 4D Nucleofector kit (Lonza, V4XC-9064). The guide RNA and Cas9 were complexed using IDT Cas9 and guides designed by Synthego. After 3-4 days, cells were expanded, and gene depletion was assessed 7 days post-electroporation via western blot analysis and/or sequencing. Guide RNA sequences can be found in Supplementary Table 9.

### PCSK9 ELISA

The concentration of PCSK9 in the conditioned media of PDAC cells was determined using the Human Proprotein Convertase 9 (PCSK9) Quantikine ELISA Kit from R&D Systems (Catalog No. DPC900), following the manufacturer’s instructions. Briefly, conditioned media from human PDAC cells were collected 24 hrs after media change, centrifuged at low speed to remove cellular debris and used in the ELISA assay.

### Flow cytometry

Cells were detached from culture vessels, resuspended in ice-cold PBS and incubated with APCconjugated anti-LDL receptor antibody (Abcam, Catalog No. ab275614) at a concentration of 0.1mg/ml for 30 mins on ice, protected from light. Following antibody incubation, cells were washed with PBS to remove unbound antibody. After the final wash, cells were resuspended in PBS for flow cytometry analysis. Stained cells were analyzed using a FACS Fortessa flow cytometer (BD Biosciences) equipped with appropriate lasers and filters for APC detection. Data were acquired using BD LSR Fortessa software and analyzed using FlowJow. Appropriate gating strategies were applied to exclude debris and ensure accurate analysis of single cells. Data were expressed as mean fluorescence intensity (MFI).

### Metabolomics

Cell pellets were collected and suspended in 100 µL of methanol in Eppendorf tubes. To each tube, 250 µL of 80% methanol and 100 μL of chloroform were added. The samples were homogenized on a MM 400 mill mixer with the aid of two 3-mm metal balls and at a shaking frequency of 30 Hz for 2 min. The homogenization step was repeated two more times. The samples were then ultra-sonicated in an ice-water bath for 2 min and subsequently centrifuged at 21,000 g and 5 °C for 10 min. The clear supernatants were collected for the following LC-MS analysis and the precipitated pellets were used for protein assay by UV-VIS spectroscopy using a standardized Bradford procedure.

For assay of sterols, 100 μL of the supernatant from each sample was mixed with 200 µL of 50% methanol, 50 µL of cholesterol-13C3 internal standard solution and 250 µL of chloroform. After vortex mixing and centrifugal clarification, the organic phase was collected and dried under a gentle flow of nitrogen gas. The residues were dissolved in 50 µL of methanol-chloroform (1:1). A solution of standard substance of sterols including cholesterol was prepared and then serially diluted with the same internal standard solution to have 9-point calibration solutions. 10-μL aliquots of the sample solutions and calibration solutions were injected to run LC-multiple-reaction monitoring (MRM)/MS on a Water Acquity UPLC system coupled to a Sciex QTRAP 6500 Plus mass spectrometer with positive-ion chemical ionization. A C8 UPLC column (2.1*50 mm, 1.7 µm) and a mobile phase of 0.1% formic acid in water (A) and 0.1% formic acid in isopropanol-acetonitrile (2:1) (B) for binary-solvent gradient elution (50% to 100% B in 18 min) at 0.4 mL/min and 60 °C were used for the chromatographic separation. The LC-MRM/MS data were acquired using Sciex Analyst 1.72 software and processed using Sciex MultiQuant 1.2 software. The linear-regression calibration curve of cholesterol was constructed with the data acquired from the calibration solutions. Concentrations of cholesterol detected in the samples were calculated by interpolating the calibration curve with the analyte-to-internal standard peak area ratios measured from the sample solutions.

### In vivo experiments

To evaluate organ seeding and growth of human PDAC cell lines, 500 000 luciferase-positive cells (KP4, Panc-1, HPAC and HPAF-II) were injected into NOD-*scid*-IL2Rgamma^null^ (NSG) mice (Jackson labs, Cat. Number 005557) via intra-splenic or tail vein injection to target the liver and the lungs, respectively. Bioluminescence imaging following administered of 30 mg/kg luciferin via intraperitoneal injection was performed weekly and imaged using an IVIS Spectrum over the duration of four weeks to monitor tumor development and progression. To measure spontaneous seeding and colonization, the same number of luciferase positive cells were injected via intracardiac injection into the left ventricle of NSG mice and imaging was performed weekly as described. Images were quantified using the “living image” software. At the end point of the experiments, tissue samples were collected and immediately fixed in 10% formalin. Following fixation, tissues embedded in paraffin and 5 micron sections were cut and subjected to H&E staining. Tumour regions within the stained tissue sections were identified based on morphological features. The percentage tumor invasion for each organ was calculated as a percentage of the total tissue area.

For intra-cardiac injection experiments, tumor infiltration in the lung and liver from the same mice was calculated as a colonisation ratio using this formula :

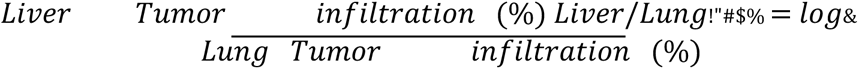

To evaluate PCSK9 abundance after organ seeding, 100 000 luciferase positive KPC derived cells (MT23 and KPC1-4), were either injected via intra-splenic, tail vein and orthotopically to target the liver, the lungs and the pancreas respectively into C57BL/6J (Black 6) mice (Jackson lab, Cat. Number 000664). Bioluminescence imaging following administered of 30 mg/kg luciferin via intraperitoneal injection was performed weekly using an IVIS Spectrum for two weeks to monitor the development and progression of tumors. At the end point of the experiments, tissue samples were collected and immediately fixed in 10% formalin. Following fixation, tissues embedded in paraffin and 5 micron sections. Tumour regions within the stained tissue sections were identified using CK19 antibody and PCSK9 abundance was evaluated.

### Mouse hepatocyte isolation and culture

Hepatocyte isolation was performed as previously described^54^. Briefly, C5BL/6 mice were anesthetized and midline laparotomy was performed to expose the liver. The vena cava was cannulated with a 27 gauge needle connected to a perfusion system. The liver was perfused sequentially with chelation buffer (EDTA-containing) to remove blood and calcium ions, followed by digestion buffer (containing liberase) to disrupt tissue structure and release hepatocytes. The liver sac was carefully opened, and the perfusate containing dissociated hepatocytes was collected in a sterile container. The collected solution was centrifuged to pellet the hepatocytes. The pellet was then resuspended in a Percoll solution and layered onto a Percoll density gradient. Centrifugation was performed to separate living hepatocytes from debris and non-parenchymal cells. Purified hepatocytes were resuspended in plating media (DMEM low glucose supplemented with 5% FBS and 1% penicillin-streptomycin) and seeded onto culture plates coated with 0.1µg/ml of collagen (Corning, Cat. Number 354236). Cells were incubated at 37°C with 5% CO2 for 3 hrs to allow initial attachment, after which plating media was replaced with maintenance media (Williams E media supplemented with 2 mM L-glutamine and 1% penicillin-streptomycin) overnight. Hepatocytes were then treated with conditioned media obtained from human PDAC cell lines as outlined in the figures. To rescue CYP46A1 KO effect, 24-hydroxycholesterol (Sigma, Cat. Number SML-1648) was used at a concentration of 25µM for the duration of the experiment (24 hours).

### Human samples

Archival de-identified sections of unmatched human primary, liver metastases and lung metastases specimens were obtained from UCSF Pathology. Human PDAC specimens were selected under Institutional review Board (IRB)-approved protocol 18-25787. PDAC patients with isolated lung metastases (M1-PUL cohort, n=10) were restrospectively identified at the Ludwig-Maximilians Uniersity of Munich, Germany, by analyzing medical records and correlating computed tomorgraphy (CT). The occurrence of M1-PUL was confirmed by histology or retrospective review of serial CT scans, which showed enlarging pulmonary nodules over time. To rule out synchronous extrapulmonary dissemination, abdominal CT scans were reviewed for the presence of extrapulmonary metastases. PDAC patients with isolated liver metastases (M1-LIV cohort, n=10) were used as a control cohort from the Ludwig-Maximilians-University of Munich, Germany. These patients were clinic-pathologically matched, as described previously^55^. FFPE blocks of these primary tumors were collected, as were patient and tumor characteristics, including age, sex, tumor-node-metastasis (TNM) stage, grading and medical information about diabetes, smoking, hypercholesterolemia, and statin usage. Project approval was granted by the ethics committee at Ludwig-Maximilians-University of Munich, Germany (approval number 134-15,20-1085, and 28410). The H&E stained sections of human lesions were examined in a blinded manner by pathologists to assess tissue morphology, cellular characteristics, and tumor features such as differentiation.

### MetMap analysis of PDAC cell line metastatic potential and penetrance

952 Data corresponding to MetMap500 was downloaded from www.depmap.org/metmap/visapp/index.html. MetMap500 provides both the metastatic potential (expressed as log10 ‘mean’ value ranging from −4 to 4) and metastatic penetrance (expressed as a value between 0-1) for 498 human cancer cell lines following intracardiac injection and subsequent seeding in 5 metastatic organs (kidney, bone, lungs, brain, or liver). For each metastatic organ site the ‘mean’ values corresponding to the 30 PDAC cell lines used in the study was transformed to the original data. The relative metastatic potential (Met^pot^) and penetrance (Met^pen^) for each organ was calculated using the formula:

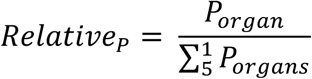

where P is either penetrance or potential, ‘organ’ refers to either kidney, bone, lungs, brain or liver and ‘organs’ refers to all organs combined. Cell lines harbouring the same potential for each organ were excluded (n = 5 cell lines). Data is graphically represented as petal plots with the length being the relative potential and color indicating penetrance.

### Principal Component Analysis (PCA)

PCA was conducted using the “fvizcluster” function from the “factoextra” package in R version.3.1093 along with the “pca_transform” function from the same package to explore the underlying structure of the metastatic potential and penetrance of the 25 PDAC cell lines. This analysis utilized the 5 organ Met^pen^ and 5 organ Met^pot^ values for each PDAC cell line. Scatter plots of the second and third principal components were generated, with points coloured according to sample groups or conditions. Petal plots were generated using the values generated to form the PCA analysis. Graphical representation was derived using GGPlot2 in R version.3.1093.

### GSEA

Gene Set Enrichment Analysis (GSEA) was performed to compare the liver and lung clusters of cell lines using gene sets derived from the signatures of Adams et al^56^ and Moffit et al^21^. The analysis was conducted using version 4.1.0 of the GSEA software. Shortly, expression data for all cell lines within the liver and lung clusters were collected and pre-processed. The gene expression profiles were normalized and log-transformed to ensure comparability across samples. Enrichment scores were calculated for each gene set, indicating the degree of enrichment in the liver cluster compared to the lung cluster, and vice versa. Statistical significance of the enrichment scores was assessed using the GSEA software.

### Correlation between liver/lung ratio and classical score

A scatter plot was generated to visualize the relationship between the liver/lung ratio and classical score derived from Adams et al.^56^ and Moffit et al.^21^ for each cell line. The liver/lung ratio was calculated by dividing the liver metastatic potential by the lung metastatic potential for each PDAC cell line according to the formula:

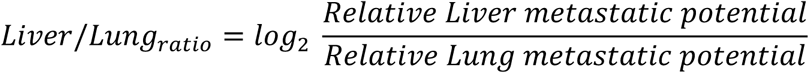

For the calculation of the classical score, we compiled all genes included in the classical gene signature used in Adams et al^56^ and Moffit et al^21^. Using gene expression data extracted from the Metmap dataset, a Z-score was computed for each classical signature associated gene for all PDAC cell lines. Aggregate Z-scores were then compiled, yielding a ‘classical score’ for each PDAC cell line. Pearson correlation coefficient (R) and p-value were calculated to assess the relationship between the liver/lung ratio and ‘classical score’.

### Gene expression correlation analysis

The DepMap RNA sequencing expression dataset was used to determine the top correlated genes associated with classical genes (GATA6, S100P, FOXQ1, CDH1) and basal genes (VIM, ZEB1). Pearson Correlation values for positively and negatively associated genes were plotted against gene rank. Red points on the scatter plot correspond to statistically significant correlations. The relative correlation of the above classical and basal genes with PCSK9 was also determined using DepMap RNA sequencing expression data. The Pearson correlation coefficient (R) and p-value for each correlation was calculated to assess the linear relationship between the two variables.

### Single cell RNA seq analysis

Datasets were uploaded from Zhang et al.^26^ and analysed as follow. Cells with nFeature_RNA being less than 200 and the percent of mitochondrial reads being more than 20 % were filtered out from the scRNA-seq datasets using the Seurat package(version 4.3.0)^57^. DoubletFinder package(version 2.0.4)^58^ was applied to find doublet cells and filter out them. Filtered data were then log normalized, scaled and integrated. The SCT assay was used to do the clustering on each sample and the integrated dataset. Cell annotation was carried out using the SingleR^59^ and the vamForSeurat function incorporating some known marker genes. InferCNV package (version 1.14.2)^60^ was implemented on each sample to identify malignant cells, designating immune cells as the reference group. Based on the inferCNV results, CNV-score and CNV-correlation were calculated. Cells exhibiting a CNV-score greater than 0.015 and a CNV-correlation exceeding 0.4 were identified as malignant cells while those with scores below these thresholds were identified as not malignant cells. Based on the cell type annotation and identified malignant cells, the RNA expression of genes PCSK9 was compared among different cell groups

### Image processing and analysis

To assess fluorescence intensity and colocalization, image analysis was conducted using Fiji software. This allowed for precise quantification of fluorescence signals and determination of colocalized areas within the images. Images acquired for analysis underwent processing utilizing Fiji software plug-ins. For crystal violet proliferation quantification, the plugin ReadPlate was employed. Fiji with automated counting was used to quantify the number of colonies present in each well.

### Statistical analysis

Statistical analysis was conducted to assess the significance of observed differences between experimental groups. For comparisons between two groups, a two-tailed t-test was employed, utilizing Prism 8 software. For experiments involving more than two groups, one-way analysis of variance (ANOVA) was utilized. Post-hoc analysis was performed to compare individual groups following ANOVA. Specifically, the Bonferroni posthoc test was employed in all cases except in vivo experiments where the Least Significant Difference (LSD) posthoc test was utilized to adjust for multiple comparisons. All statistical analyses were performed using standard statistical software. packages. Statistical significance was defined as p < 0.05. Pearson correlation were performed using the ggpubr package (version 0.6.0), the stat_cor function with “pearson” method

